# The heteromeric *Plasmodium falciparum* pantothenate kinase has only one active site and does not require *Pf*14-3-3I for activity

**DOI:** 10.64898/2026.01.22.701186

**Authors:** Xiangning Liu, Christina Spry, Kevin J Saliba

## Abstract

Coenzyme A (CoA) is an essential molecule for the intraerythrocytic stage of *Plasmodium falciparum*. Pantothenate kinase (PanK) catalyses the first step of the CoA biosynthesis pathway and functions as a homodimer in most organisms investigated thus far. *P. falciparum* possesses a novel heteromeric PanK complex composed of *Pf*PanK1, *Pf*PanK2 and *Pf*14-3-3I. Using a mutagenesis approach, we generated 10 *Pf*PanK mutants and demonstrate that the *Pf*PanK complex has only one functional active site, with both *Pf*PanK1 and *Pf*PanK2 required for activity by the complex. We also show that *Pf*PanK2 is essential for normal intraerythrocytic parasite proliferation using a conditional knockdown system. 14-3-3 binding motifs generally contain a phosphoserine/threonine residue. Mass spectrometry analyses of phospho-peptide enriched, immunoprecipitated *Pf*PanK samples revealed phosphorylation sites in both *Pf*PanK1 and *Pf*PanK2 that were additional to the previously reported sites. To investigate the role of specific sites in *Pf*PanK1 and *Pf*PanK2 that may be involved in *Pf*14-3-3I binding, five additional mutants were generated. Mutagenesis of four predicted *Pf*14-3-3I binding sites in *Pf*PanK1 resulted in a significant reduction in the amount of *Pf*14-3-3I bound to the *Pf*PanK complex, with S334 being the most likely binding site. Heterologous expression of the *Pf*PanK complex in an insect cell system yielded a small amount of soluble protein that assembled *in situ* into a functional complex. Combined results from heterologous expression and *P. falciparum* mutagenesis suggest that *Pf*14-3-3I may not be essential for *Pf*PanK activity but may be important for stabilising the *Pf*PanK complex.

## Introduction

Malaria is a lethal disease that affects tropical and subtropical countries. It is spread by mosquitoes and caused by parasites of the genus *Plasmodium*. There were an estimated 263 million new cases in 2023 resulting in approximately 597,000 deaths worldwide (1). Despite remarkable progress towards malaria elimination, the emergence and spread of parasites resistant/partial resistant to artemisinin combination therapies (ACTs)(2–5), the current frontline treatment for malaria, is a major concern. Therefore, there is an urgent need for novel antimalarials with unique mechanisms of action to enter the developmental pipeline so that we can continue to have safe treatments for malaria (6, 7).

Vitamin B_5_ (pantothenate) is an essential water-soluble vitamin required by the intraerythrocytic stage parasite. It is the precursor to coenzyme A (CoA), which is an essential enzyme factor that functions as an acyl group carrier and activator in key cellular processes in all living organisms (8–10). Since the discovery of the antimicrobial activity of pantothenate derivatives in the 1940s, the CoA biosynthesis pathway in *Plasmodium falciparum*, the deadliest *Plasmodium* species that infects humans, has received considerable attention as a potential drug target (11–16).

Pantothenate kinase (PanK) is the first enzyme in the CoA biosynthesis pathway. It catalyses the ATP-dependent phosphorylation of pantothenate to 4’-phosphopantothenate. In many organisms, this step determines the rate of CoA biosynthesis as it is subject to feedback inhibition (9, 17, 18). In *P. falciparum*, the phosphorylation of pantothenate by PanK(s) in parasite lysate has also been shown to be subjected to CoA-mediated feedback inhibition (19, 20).

Three phylogenetically distinct types of PanK (type I, type II or type III) have been characterised, delineated by their differences in structures, catalytic properties and inhibition profiles (10, 21). Type I PanKs are found exclusively in bacteria, type II PanKs are found mainly in eukaryotes, while type III PanKs are found in many pathogenic bacteria either alone or in addition to another PanK type (21–23). Many organisms encode multiple PanKs, *i.e.* some express two different PanK types, while others carry two different PanKs of the same type. Although a common phenomenon, the functional significance of cells possessing multiple PanKs is poorly understood. To date, all PanKs with a solved structure exist as homodimers (24–31).

Apicomplexans typically possess two type II PanKs (referred to PanK1 and PanK2) (32, 33). Tjhin *et al.* (33) have shown that *P. falciparum* and *Toxoplasma gondii* harbor a novel heteromeric PanK complex, making it unique from other homodimeric bacterial and mammalian PanKs reported. Consistent with the two PanKs not being functionally redundant, a genome-wide insertional mutagenesis study (34) has deemed both *P. falciparum* PanKs (*Pf*PanKs) to be important for the optimal proliferation of intraerythrocytic stage of the *P. falciparum* parasite. Tjhin *et al.* (20) have also shown that mutations in *Pf*PanK1 alter PanK activity in *P. falciparum* and impair parasite proliferation. *Pf*PanK2 is less well studied in *P. falciparum*, however, in *T. gondii*, *Tg*PanK2 is essential for PanK function and parasite proliferation (33). By comparing the amino acid sequences of *P*. *falciparum* and *T*. *gondii* PanKs with those of other type II PanKs, it was proposed that PanK homodimers from *P. falciparum* and *T. gondii* are likely not functional (if they even form), and that only heteromeric PanK1/PanK2 complexes in these parasites, can serve as functional PanKs (33).

The heteromeric *Pf*PanK complex contains a candidate regulatory protein termed *Pf*14-3-3I (33). It is a 30 kDa protein which belongs to a family of conserved multitask molecules that are widely distributed among eukaryotes and have a variety of regulatory functions (e.g. cell cycle regulation, signal transduction and apoptosis) (35–37). These functions are carried out by 14-3-3 proteins by modifying the trafficking/targeting (38), conformation, localisation, and/or activity of their target protein (39). Often existing as either homo- or heterodimers, 14-3-3 proteins form extremely stable complexes. The rigidity of the 14-3-3 dimer can force structural rearrangements on their target proteins, regulating their activity and properties (40, 41). Although it has been shown that *Pf*PanK1 and *Pf*PanK2 function in a heteromeric complex that involves *Pf*14-3-3I, the exact role of *Pf*14-3-3I and its binding site remain to be elucidated.

In this study, we demonstrate that the heteromeric *Pf*PanK complex has a single complete active site, unlike PanKs in any other organism reported to date. We show for the first time, that *Pf*PanK2 contributes to PanK function and is essential for the optimal growth of intraerythrocytic stage of *P. falciparum*. Furthermore, we have identified novel phosphorylation sites in *Pf*PanK1 and *Pf*PanK2 and the potential binding site(s) of *Pf*14-3-3I in *Pf*PanK1. We have also heterologously expressed the *Pf*PanK complex in insect cells. We show that the heteromeric complex assembles *in situ*, is functional and unlikely to require *Pf*14-3-3I for assembly or activity.

## Results

### *Pf*PanK1 and *Pf*PanK2 associate into the same complex endogenously

To confirm that the previously-described (33) heteromeric PanK complex is of physiological relevance in *P. falciparum*, we generated transgenic parasites expressing a GFP-tagged *Pf*PanK1 or *Pf*PanK2 under control of the endogenous promoter. We first examined the sub-cellular localisation of endogenously GPF-tagged *Pf*PanK1 and *Pf*PanK2. In both cases, the fluorescence was observed throughout the parasite cytosol, with the regions of higher fluorescence intensity overlapping with the nucleic acid stain (**Figure 1A**). These observations are consistent with the previous data on episomally-expressed *Pf*PanK-GFP fusion proteins (20, 33). On the denaturing western blot, a band at the expected molecular weight for *Pf*PanK1-GFP (∼87 kDa) or *Pf*PanK2-GFP (∼117 kDa) was detected, confirming successful expression of the endogenous fusion proteins (**Figure 1B**). Native western blotting and fluorescence-coupled size exclusion chromatography (FSEC) were then performed and the heteromeric PanK complex could be detected using both approaches (**Figure 1C and 1D**), confirming the presence of this complex when expressed from the endogenous locus in *P. falciparum*. Consistent with the previous report of Tjhin *et al.* (33), *Pf*14-3-3I was found to be part of the *Pf*PanK complex when expressed from the endogenous locus (**Figure S1**).

**Figure 1.**
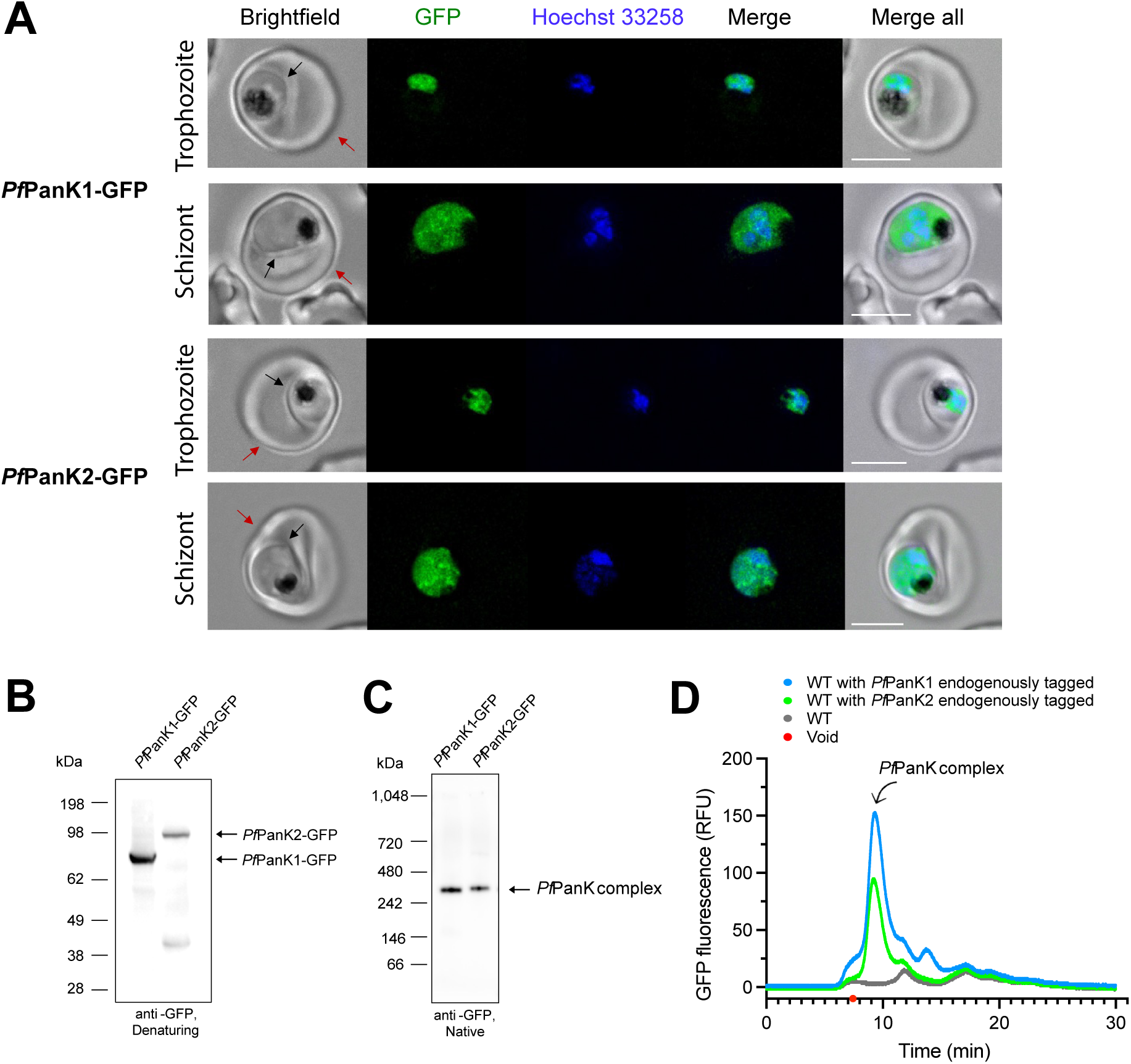
Endogenous *Pf*PanK complex in *P. falciparum* parasites. **(A)** Fluorescence microscopy of trophozoite or schizont stage parasites endogenously expressing *Pf*PanK1-GFP or *Pf*PanK2-GFP. Nuclei of parasites were stained with Hoechst 33258. From left to right: brightfield; GFP fluorescence (green); Hoechst 33258 (blue); merge of the two fluorescent channels; merge of all three channels. The parasite membrane is indicated with a black arrow and the erythrocyte membrane is indicated with a red arrow. Scale bar: 5 µm. **(B)** Denaturing western blots using anti-GFP antibody showing the expression of *Pf*PanK1-GFP and *Pf*PanK2-GFP: *Pf*PanK1 (left) and *Pf*PanK2 (right). The expected sizes of the proteins are ∼87 kDa for *Pf*PanK1-GFP and ∼117 kDa for *Pf*PanK2-GFP. **(C)** Anti-GFP native western blot showing the *Pf*PanK1-GFP (left) and the *Pf*PanK2-GFP (right) complexes in parasite lysates. **(D)** Fluorescence-coupled size exclusion chromatography traces of GFP-tagged proteins from *Pf*PanK1-GFP, *Pf*PanK2-GFP and wild-type (WT) parasites. The endogenous *Pf*PanK complex was observed in both transgenic parasite lines and was absent in the non-tagged WT parasites. Void is marked with a red dot on the x-axis.

### The *Pf*PanK complex possesses a single complete active site

To investigate why *Pf*PanKs exist in a heteromeric complex, Tjhin *et al.* (33) compared the amino acid sequences of the *Pf*PanKs with that of *Hs*PanK3, a well-studied type II PanK (Tjhin *et al.*, 2021, **Figure 2A**). In the homodimeric *Hs*PanK3, residues from both protomers contribute to the stabilisation of the two identical active sites and their interaction with pantothenate. E138 from protomer 1 forms a hydrogen bond with Y254 from protomer 2 and D137 from protomer 1 with Y258 from protomer 2 (**Figure 2A**, (42)). Multiple sequence alignment analyses revealed that *Pf*PanK1 and *Pf*PanK2 do not individually contain the complete set of residues required for active site stabilisation. For instance, D137 is only conserved in *Pf*PanK1 while the two tyrosines are only conserved in *Pf*PanK2. It is therefore hypothesised that homodimeric *Pf*PanK1 or *Pf*PanK2 are unlikely to be functional, and that only a heteromeric *Pf*PanK1+*Pf*PanK2 complex, with a single complete active site, can act as a functional PanK (33).

**Figure 2.**
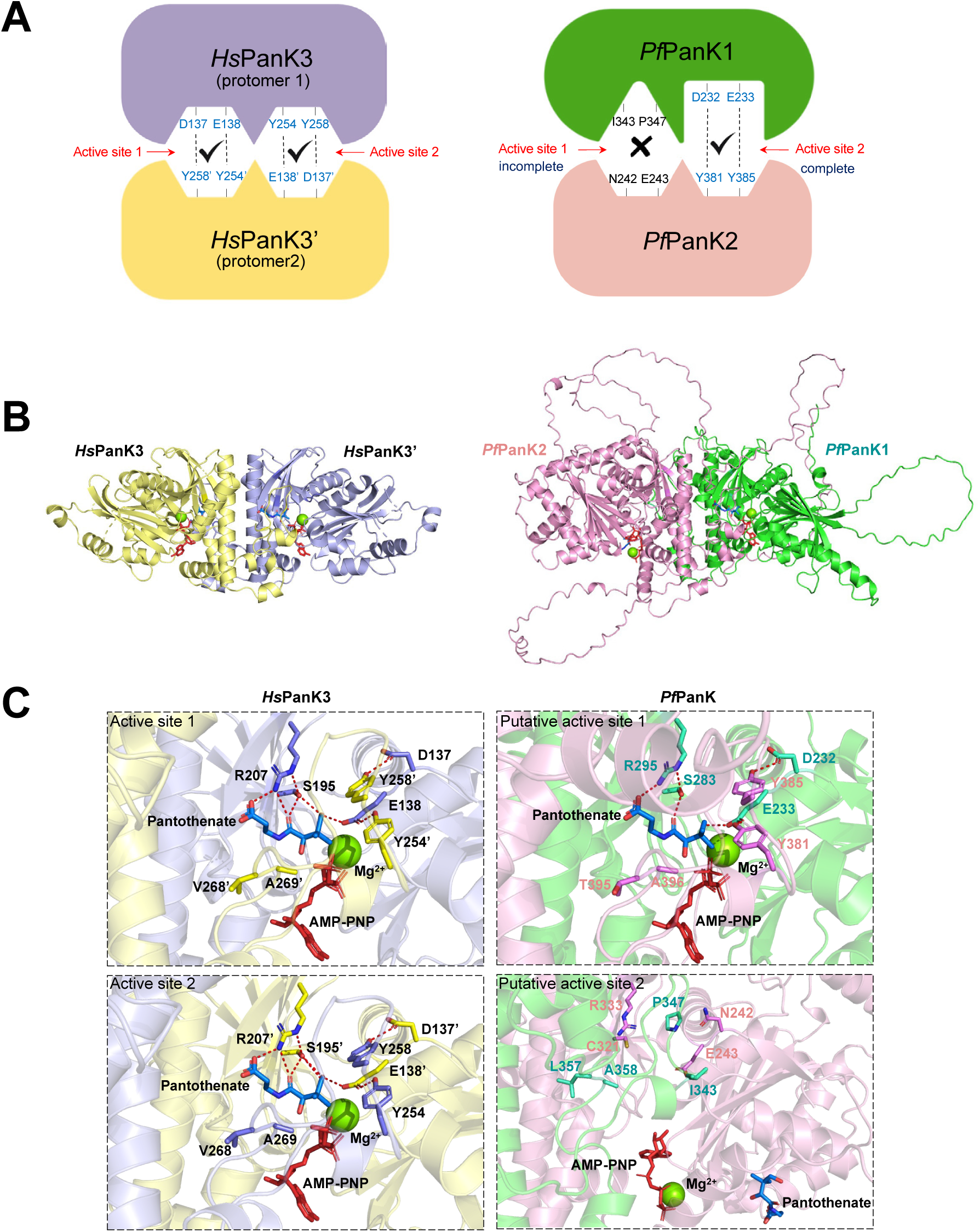
Active sites in *Hs*PanK3 homodimer and *Pf*PanK heterodimer. **(A)** A cartoon showing that the homodimeric *Hs*PanK3 is composed of two identical protomers (yellow) forming two identical active sites. Residues from both protomers participate in the stabilisation of the active site and interact with pantothenate (42). The heteromeric *Pf*PanK is composed of *Pf*PanK1 (green) and *Pf*PanK2 (pink). Residues from both *Pf*PanK1 and *Pf*PanK2 contribute to the stabilisation of the active site in *Pf*PanK. Hydrogen bonds between residues are in dashed lines. An apostrophe denotes residues from protomer 2 in *Hs*PanK3. The cartoons are based on a crystal structure for *Hs*PanK3 and a model of equivalent residues for *Pf*PanK. **(B)** Crystal structure of the AMP-PNP-pantothenate-bound state of homodimeric *Hs*PanK3 (PDB code: 5KPR, left) and the AlphaFold predicted heterodimeric *Pf*PanK1+*Pf*PanK2 complex with AMP-PNP and pantothenate docked in (right). **(C)** Magnification of the two active sites in homodimeric *Hs*PanK3 and heterodimeric *Pf*PanK. In the homodimeric *Hs*PanK3 (left), residues from both protomers contribute to the stabilisation of the binding pocket (E138 forms a hydrogen bond with Y254’ and D137 with Y258’) and interact with pantothenate (E138, S195, R207, A269’ and V268’). In the heterodimeric *Pf*PanK complex, the two active sites formed are non-identical (right). The ‘complete’ active site formed (top right) closely resembles the two active sites in *Hs*PanK3. Hydrogen bonds are formed between D232 (*Pf*PanK1) and Y381(*Pf*PanK2), E233 (*Pf*PanK1) and Y385 (*Pf*PanK2), which stabilise the active site. In contrast, the ‘incomplete’ site formed (bottom right) does not contain the two sets of hydrogen bonds required for the stabilisation of the active site. AMP-PNP and pantothenate could not be docked appropriately in the incomplete site in any of top five docking poses. Hydrogen bonds with and between the sidechains of these residues are shown with red dashed lines. An apostrophe denotes residues from the lilac-coloured protomer in *Hs*PanK3.

To test this hypothesis, we first generated a model for the heteromeric *Pf*PanK complex using AlphaFold (**Figure 2B**). We did not include *Pf*14-3-3I in this model as the interaction of *Pf*PanK1 and *Pf*PanK2 with the amphipathic binding grove of *Pf*14-3-3I was predicted with very low confidence (per-residue Local Distance Difference Test, pLDDT <50). The dimerisation surface, which is composed of the C-terminal ⍺9 helix in *Hs*PanK3, is conserved in both *Pf*PanKs. It is worth noting that, despite *Pf*PanK2 containing an additional 268 amino acid insert in the loop that has been associated with dimer formation (29, 32), its ability to dimerise with *Pf*PanK1 is not affected, as evidenced by the presence of the complex when expressed from the endogenous locus (**Figure 1B and 1C**). Two non-identical active sites in the *Pf*PanK complex are predicted with high confidence (pLDDT >70 but <90).

To gain better insights into the catalytic mechanism of the *Pf*PanK complex, we performed molecular docking to place pantothenate, adenylyl-imidodiphosphate (AMP-PNP, the non-hydrolysable ATP analogue) and magnesium ion into the two active sites. As shown in **Figure 2C**, in contrast to homodimeric *Hs*PanK3, which possess two identical active sites (left panel), the two sites formed by heterodimeric *Pf*PanK1+*Pf*PanK2 complex are different (right panel). The docking results are consistent with the multiple sequence alignment results and the hypothesis that the heteromeric *Pf*PanK complex only possesses a single complete active site. Similar to *Hs*PanK3 (**Figure 2C, left panel**), *Pf*PanK1 and *Pf*PanK2 residues predicted to form and stabilise the complete active site, adopt comparable orientations (**Figure 2C**, **top right panel**). Hydrogen bonds are formed between D232 (*Pf*PanK1) and Y381 (*Pf*PanK2), E233 (*Pf*PanK1) and Y385 (*Pf*PanK2) as predicted, reflecting the interactions of the corresponding residues in *Hs*PanK3 that are vital for active site stabilisation. (42). Pantothenate, AMP-PNP and a magnesium ion were successfully docked into the complete active site (AMP-PNP: - 10.355 kcal/mol, pantothenate: -6.834 kcal/mol) and were orientated in the active site in the same way as observed in the AMP-PNP-pantothenate-bound *Hs*PanK3 crystal structure. E233, S283, R295 (from *Pf*PanK1), T395, A396 (from *Pf*PanK2) interact with pantothenate. Anchoring of the carboxyl group of pantothenate in the active site is achieved by R295 of *Pf*PanK1 as expected, which is further facilitated by S283 of *Pf*PanK1 by hydrogen bonding.

The apparently ‘incomplete’ site formed by *Pf*PanK1 and *Pf*PanK2 is different to the complete site (**Figure 2C, bottom right panel**). The corresponding residues to Y254 and Y258 (*Hs*PanK3 numbering) in *Pf*PanK1 are I343 and P347, respectively, which are unable to form hydrogen bonds with N242 and E243 of *Pf*PanK2 and therefore cannot stabilise the site. The ‘incomplete’ nature of this site is further supported by our inability to dock pantothenate and AMP-PNP into this site. Taken together, our AlphaFold-predicted model of the *Pf*PanK1+*Pf*PanK2 complex and the subsequent molecular docking results are consistent with the hypothesis that the heteromeric *Pf*PanK complex only possesses a single complete active site.

To confirm our Alpha-Fold results, we performed site directed mutagenesis of five residues (D232, E233, R295 in *Pf*PanK1 and Y381, Y385 in *Pf*PanK2) predicted to be important for active site stabilisation in the complete active site and the corresponding ones (I343, P347 in *Pf*PanK1 and N242, E243, R333 in *Pf*PanK2) in the incomplete site. The mutated proteins were GFP-tagged and episomally overexpressed in *P. falciparum*. Using fluorescence microscopy, all 10 mutant *Pf*PanK1-GFP / *Pf*PanK2-GFP proteins were found to localise to the parasite cytosol and not be excluded from the nucleus, as observed for the WT proteins, suggesting that none of the mutations affected localisation (**Figure S2** and **S3**). To examine any impact of mutation on the formation of the *Pf*PanK complex, proteins extracted from each mutant parasite were analysed by western blots under non-denaturing native conditions. The *Pf*PanK complex band was observed in all *Pf*PanK1 and *Pf*PanK2 mutants, consistent with these mutations not influencing the assembly of the *Pf*PanK complex (**Figure 3A**).

**Figure 3.**
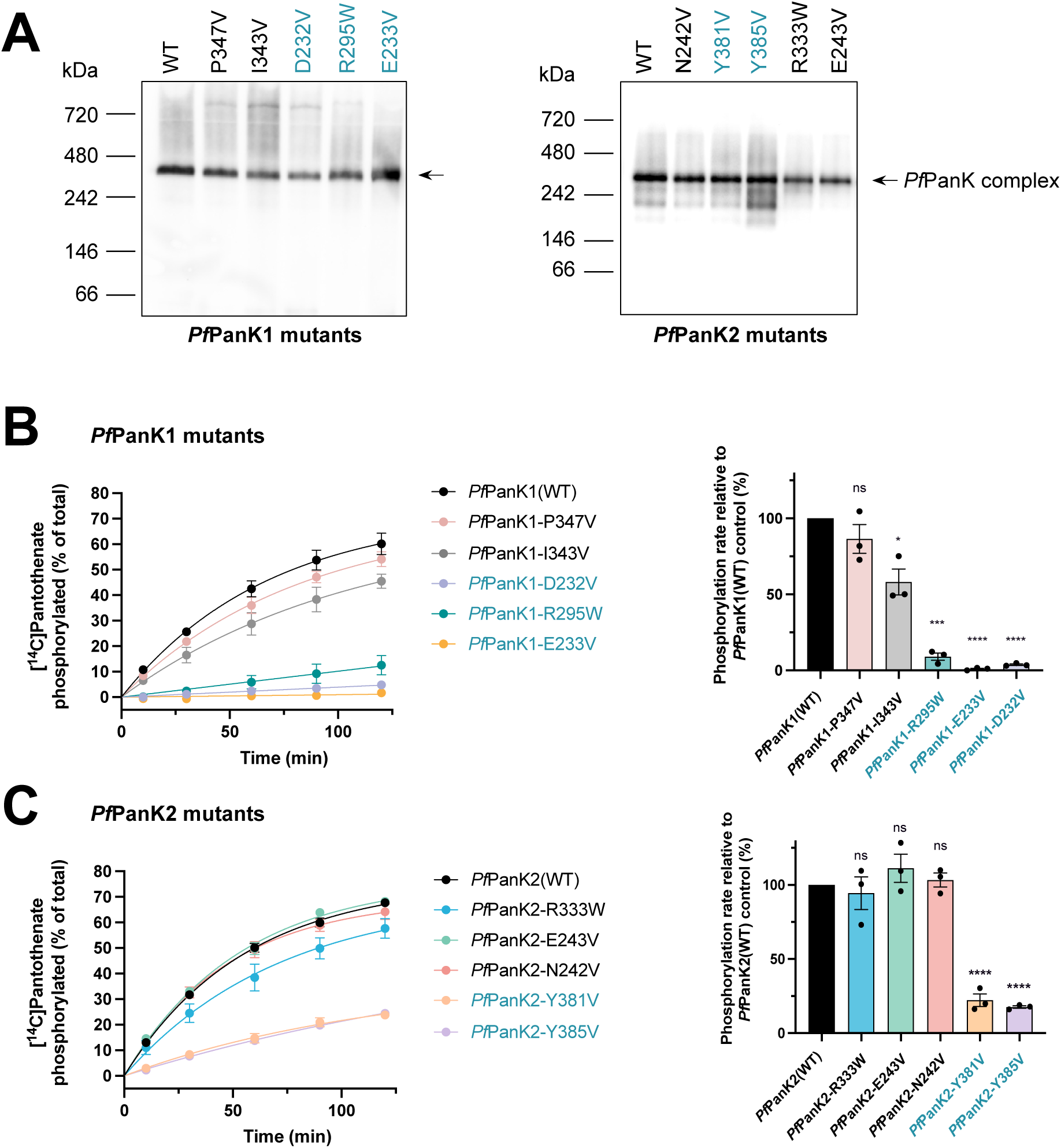
Analyses of *Pf*PanK1 and *Pf*PanK2 mutants. Text relating to residues situated in the complete site is colored tuiquoise while that relating to residues of the incomplete site are shown in black. **(A)** Anti-GFP native western blots showing the *Pf*PanK-GFP complex bands in lysates prepared from all mutant parasites, indicating that none of the introduced mutations influenced the assembly of the complex. The western blot shown is a representative of three independent experiments, each performed with a different batch of parasites. **(B)** The phosphorylation of [^14^C]pantothenate over time (left) by GFP-Trap immunoprecipitated fractions prepared from WT control *Pf*PanK1 and mutant *Pf*PanK1 transgenic parasites. Phosphorylation rates (right) were determined from the overall reaction time course using the one-phase association equation in GraphPad Prism (for *Pf*PanK1(WT), *Pf*PanK1-I343V and *Pf*PanK1-P347V) or linear regression (for *Pf*PanK1-D232V, *Pf*PanK1-E233V and *Pf*PanK1-R295W). **(C)** The phosphorylation of [^14^C]pantothenate over time (left) by GFP-Trap immunoprecipitated fractions prepared from WT control *Pf*PanK2 and mutant *Pf*PanK2 transgenic parasites. Phosphorylation rates (right) were determined from the overall reaction time course by one-phase association for all samples. Values are presented as a percentage relative to the WT control phosphorylation rate. Values are averaged from three independent experiments, each performed with a different batch of parasites and carried out in triplicate. Error bars represent SEM. Significant differences in phosphorylation rate are marked with asterisks (ns – not significantly different, * P ≤ 0.05, ** P ≤ 0.01, *** P ≤ 0.001 and **** P ≤ 0.0001).

Next, GFP-tagged mutant or WT *Pf*PanK complex was immunoprecipitated using GFP-Trap and the coimmunoprecipitated fractions were then used in a [^14^C]pantothenate-based phosphorylation assay to assess the kinetic activity of each mutant *Pf*PanK complex (**Figure S4**). Mutation of three *Pf*PanK1residues located in the predicted complete active site (D232, E233 and R295, **Figure 2A and 2D**) resulted in severely reduced PanK activity as predicted (**Figure 3B**). Mutation of D232 and E233, the active-site stabilising residues, to valine eliminated PanK activity of the complex (ANOVA, p ≤ 0.0001), while the R295W mutated complex displayed a 90% lower phosphorylation rate as compared to the WT *Pf*PanK complex (ANOVA, p = 0.0002). The equivalent *Hs*PanK3 active-site stabilising residues Y251’ and Y258’ in *Pf*PanK1 are I343 and P347, both of which are located in the predicted incomplete site of the *Pf*PanK complex (**Figure 2D**). No significant difference between the phosphorylation rate of the WT and P347V *Pf*PanK complexes was observed (ANOVA, p>0.05, **Figure 3B**). For the I343V mutated complex, we observed a 40% reduction in the phosphorylation rate (ANOVA, p = 0.016, **Figure 3B**). This is unexpected as I343 is situated in the predicted ‘incomplete’ site of the *Pf*PanK complex.

In *Pf*PanK2, mutation of R333, E243 or N242 did not influence the activity of the PanK complex as hypothesised since they are all situated in the predicted ‘incomplete’ site (ANOVA, p>0.05; **Figure 3C**). On the other hand, mutation of Y381 or Y385, which are predicted to form hydrogen bonds with D232 and E233 from *Pf*PanK1 to stabilise the complete active site (**Figure 2D**), resulted in ∼80% reduction in the phosphorylation rate compared to the WT complex (**Figure 3C**, ANOVA, p<0.0001). Overall, our site-directed mutagenesis data are consistent with the docking results and support our hypothesis that the heteromeric *Pf*PanK complex only has one complete active site, with residues from both *Pf*PanK1 and *Pf*PanK2 contributing.

### *Pf*PanK2 is essential for normal intraerythrocytic proliferation of *P*. *falciparum*

Having confirmed that the heteromeric *Pf*PanK complex possesses only one complete active site and that *Pf*PanK2 contributes to PanK activity, we envisage that *Pf*PanK2 is also important for *P. falciparum* proliferation, as has been observed in *T. gondii* (33). To begin investigating the importance of *Pf*PanK2 for proliferation, we engineered the CJ-15,801-A (CJ-A) strain *P. falciparum*, which gave rise to a transgenic parasite line referred to as *Pf*PanK2-*glmS*. In this line, the gene encoding *Pf*PanK2 is modified to incorporate a C-terminal GFP tag and the GlcN inducible *glmS* ribozyme sequence within the terminator region (**Figure 4A**). The CJ-A parasite line was generated by *in vitro* evolution using CJ-15,801-pressure and harbors a G95A mutation in *Pf*PanK1, which results in significantly reduced PanK activity (20). In a six-week competition assay, Tjhin *et al.* (20) showed that the G95A mutation results in a significant fitness cost to the parasite, consistent with *Pf*PanK1 being important for normal intraerythrocytic proliferation of *P*. *falciparum*. We therefore reasoned that *Pf*PanK2 knockdown within the CJ-A parasite background would cause a parasite growth defect in a shorter period of time when compared to *Pf*PanK2 knockdown within the wild-type background. Correct integration of the targeting knockdown construct and the clonality of the generated transgenic parasites were confirmed by diagnostic PCR (**Figure 4B**). Next, *Pf*PanK2 was knocked down by exposing the *glmS*-regulated line to 5 mM glucosamine (GlcN), the highest concentration of the inducer that we determined was not detrimental to the proliferation of the parent line (*i.e* CJ-A) over 96 h. Successful knockdown of *Pf*PanK2-GFP as well as the *Pf*PanK complex (since *Pf*PanK2 is integral to the complex) was observed (**Figure 4C**). Densitometry analyses showed no decrease in *Pf*HSP70 (the loading control) expression in both GlcN-treated and non-treated parasites over 96 h. In contrast, a ∼65% reduction in *Pf*PanK2-GFP and the *Pf*PanK complex expression was observed in GlcN-treated parasites (**Figure 4D**).

**Figure 4.**
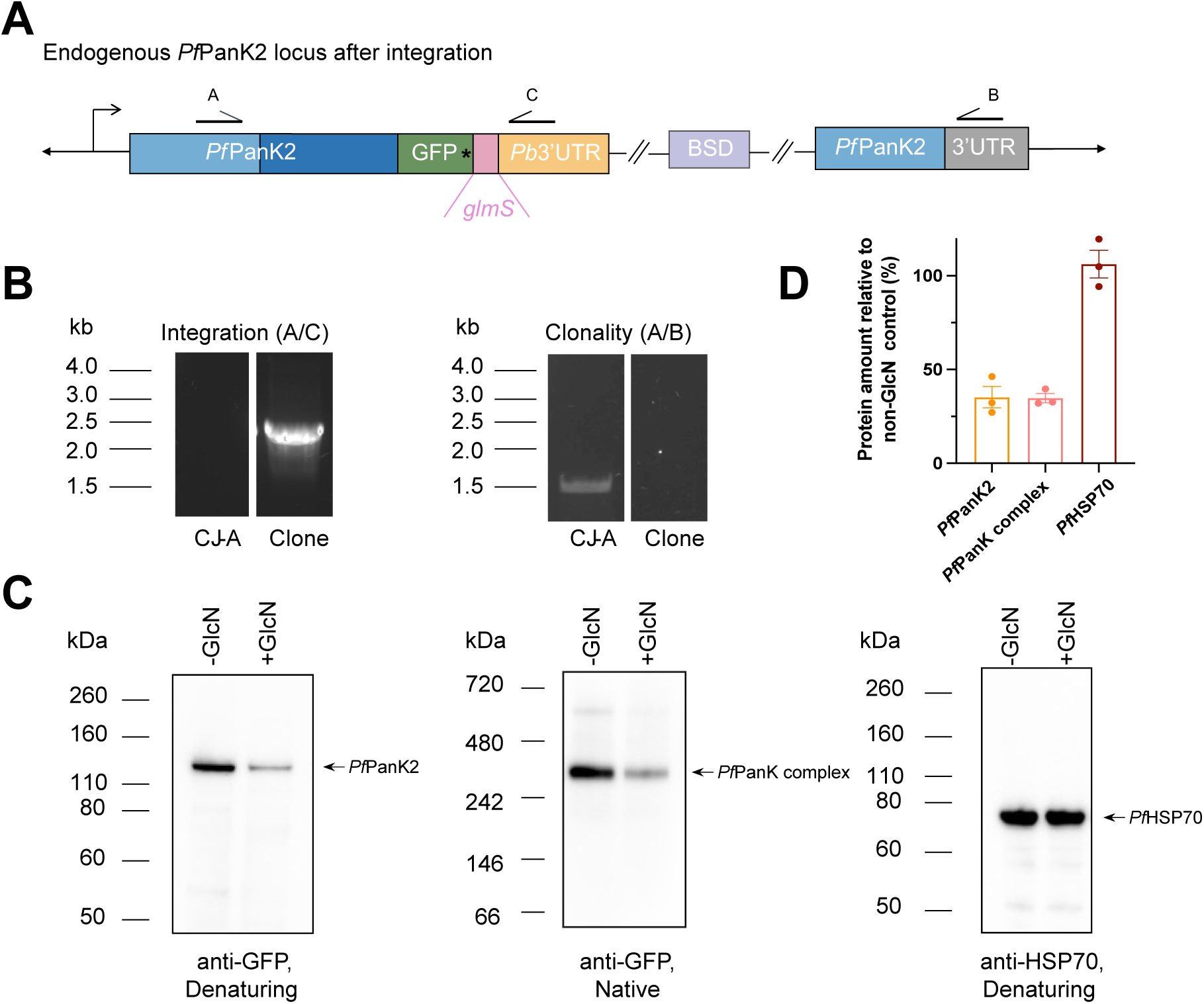
Knockdown of *Pf*PanK2 protein expression. **(A)** Schematic showing the recombinant *Pf*P*anK2* locus after incorporating the *glmS* ribozyme and GFP at the 3’ end by single crossover homologous recombination. BSD denotes the blasticidin resistance cassette and the stop codon is marked by an asterisk. The *Pb*3′UTR represents the 3′ untranslated region derived from *Plasmodium berghei*, while the 3′UTR corresponds to the endogenous 3′ untranslated region of *Pf*PanK2. **(B)** Diagnostic PCRs using parent (CJ-A) and transgenic parasite gDNA as template. Arrows on the recombinant *Pf*PanK2 locus indicate binding sites for diagnostic PCR primers. Presence of the 2494 bp product (using primer A and C) indicates the integration of the desired construct (left). Absence of the 1512 bp product (using primer A and B) indicates these transgenic parasite lines are clonal (right). **(C)** GlcN-induced knockdown of *Pf*PanK2-GFP protein at 96 h. Membranes were probed with anti-GFP antibody to detect the *Pf*PanK2-GFP protein (left) and the *Pf*PanK complex (middle), and with anti-HSP70 as a loading control (right). The western blot shown is a representative of three independent experiments, each performed with a different batch of parasites. **(D)** Densitometry analyses of the knockdown western blots. The amount of protein detected in the GlcN-treated parasites at 96 h is normalised to the amount of the protein detected in the non-treated parasites and expressed as a percentage. Values are averaged from three independent experiments and error bars represent SEM.

To determine the minimal pantothenate concentration suitable for assessing the effect of *Pf*PanK2 knockdown on parasite proliferation, we examined the pantothenate requirement of the parent and the *Pf*PanK2-*glmS* parasites in the absence of GlcN, as an extracellular supply of pantothenate is essential for the *in vitro* proliferation of the intraerythrocytic stage of *P. falciparum* (43). We have observed a subtle yet statistically significant increase (∼10 nM) in the 50% stimulatory concentration (SC_50_) of pantothenate (the concentration of pantothenate required to support parasite proliferation to a level equivalent to 50% of the control level (20)) for the *Pf*PanK2-*glmS* parasites compared to CJ-A parent parasites (unpaired two-tailed *t*-test, p = 0.0056; **Figure 5A**). It is possible that the incorporation of the GFP-tag on the C-terminus of *Pf*PanK2 might influence the stability of the *Pf*PanK complex, resulting in slightly reduced PanK activity (in the absence of GlcN) and therefore, an increase in the exogenous pantothenate requirement of *Pf*PanK2-KD parasites.

**Figure 5.**
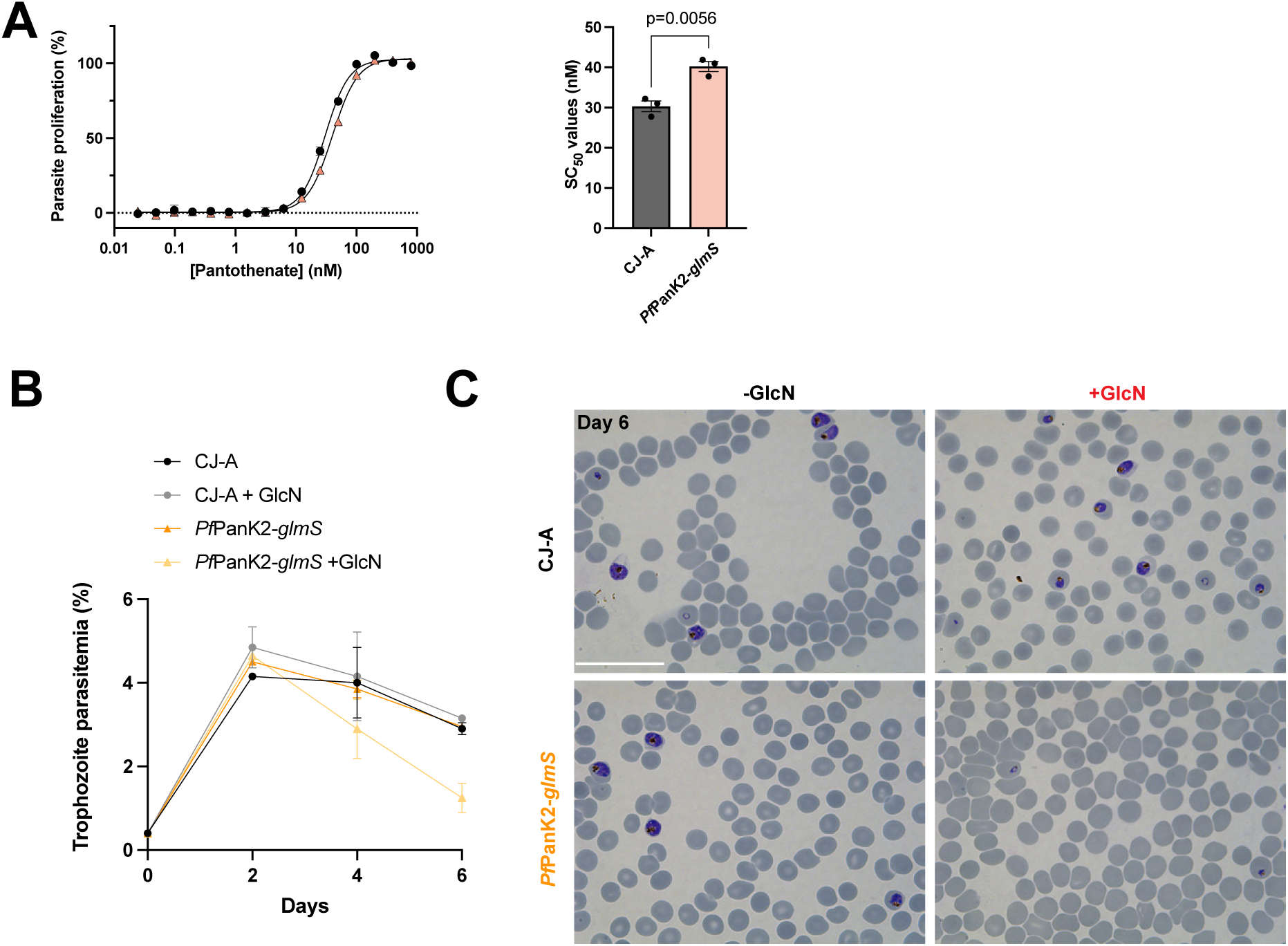
*Pf*PanK2 is essential for normal intraerythrocytic proliferation of *P. falciparum*. **(A)** Percentage proliferation of CJ-A (parent) and *Pf*PanK2-*glmS* parasites grown in different pantothenate concentrations. Values are averaged from three independent experiments, each carried out in triplicate. Error bars represent SEM and are not visible if smaller than the symbols. A statistically significant difference in pantothenate stimulatory concentration 50 (SC_50_) was observed. **(B)** In the presence of 200 nM pantothenate, growth of CJ-A and *Pf*PanK2-*glmS* parasites over three intraerythrocytic cycles in the presence or absence of GlcN. At the end of each intraerythrocytic growth cycle, parasites were harvested for FACS analysis to determine parasitemia levels. Parasites were added to fresh erythrocytes at 0.4% parasitemia on day 0, diluted 1 in 15 on day 2 and then diluted 1 in 20 on day 4 for all cultures. Values are averaged from two independent experiments and error bars represent range/2. **(C)** Light microscopy of Giemsa-stained *Pf*PanK2-*glmS* parasites (on day six) shows that the proliferation of GlcN-treated parasites in the presence of 200 nM pantothenate was impaired and substantially delayed in the intraerythrocytic cycle. No major difference was observed between GlcN-treated and non-treated CJ-A parent parasites. Scale bar represents 31.1 µm. Images are representative of 6 fields examined from 2 independent experiments.

To investigate a potential effect of *Pf*PanK2 knockdown on parasite proliferation, we exposed the *Pf*PanK2-*glmS* and CJ-A parent line parasites to 5 mM GlcN for six days in the presence of 200 nM exogenous pantothenate (the lowest concentration required to support 100% proliferation of *Pf*PanK2-KD parasites in the absence of GlcN), and compared their proliferation to control parasite cultures not treated with GlcN. For *Pf*PanK2-*glmS* parasites, trophozoite parasitaemia decreased to below 1.5% after six days in the presence of GlcN (**Figure 5B**) and the parasites were substantially delayed in the intraerythrocytic cycle (**Figure 5C**), as opposed to the non-treated parasites, which proliferated normally throughout the experiment time-course (parasitaemia of 3.0%). For the CJ-A parent parasites, no obvious difference in proliferation was observed between the GlcN-treated and non-treated parasites (**Figure 5B** and **5C**), confirming that the impaired proliferation we observed in the GlcN-treated *Pf*PanK2-*glmS* culture was due to the knockdown of *Pf*PanK2 expression. These data indicate that *Pf*PanK2 is essential for normal intraerythrocytic proliferation of *P. falciparum*, as observed previously for *Pf*PanK1 (20) and, notably, that neither can substitute for the other. Whether the same outcome would be observed if the *Pf*PanK2 KD was performed in WT parasites remains to be determined.

### Identification of *Pf*PanK phosphorylation sites and exploration of their role in binding to *Pf*14-3-3I

In addition to *Pf*PanK1 and *Pf*PanK2, *Pf*14-3-3I appears to be an integral component of the heteromeric *Pf*PanK complex (33). The role of *Pf*14-3-3I in the complex and its binding site is unknown. Binding to 14-3-3 typically requires a specific target motif with a phosphoserine/threonine residue (44). In intraerythrocytic stage parasites, no phosphorylated residues have been identified in *Pf*PanK1, but up to seven serine residues (S52, S55, S484, S516, S569, S571 and S621) have been shown to be phosphorylated in *Pf*PanK2 in five independent phospho-proteomic studies (45–49). Only two of them (S52 and S55) are within *Pf*14-3-3I binding sites predicted for *Pf*PanK2. To explore the possibility that further serine/threonine residues in *Pf*PanK1 and/or *Pf*PanK2 are also phosphorylated, and could form part of a *Pf*14-3-3I binding site, we immunoprecipitated the *Pf*PanK complex and performed phospho-peptide enrichment prior to liquid chromatography-tandem mass spectrometry (LC-MS/MS) analysis.

We detected four phosphorylated sites in *Pf*PanK1 (S11, S160, S484 and S487; highlighted turquoise or yellow in **Figure 6A**). None of these residues coincide with bioinformatically predicted *Pf*14-3-3I binding sites within *Pf*PanK1 (**Figure 6A**). It should be noted that peptide 89-<u>ITLTGG</u>GAHK-98 and 287-SNGYDSYQRIAGTAIGGGTLMGLAKIILDNISFEELIKCAE<u>DKNKNISFDLK</u>-338, each containing a predicted *Pf*14-3-3I binding site (underlined), were confidently identified as phospho-peptides. The location of the phosphorylation site(s) on these peptides, however, could not be confidently determined. It is therefore possible that T90 and S334 are phosphorylated (T92 is not predicted to be required for 14-3-3 binding) and could form part of the *Pf*14-3-3I binding site.

**Figure 6.**
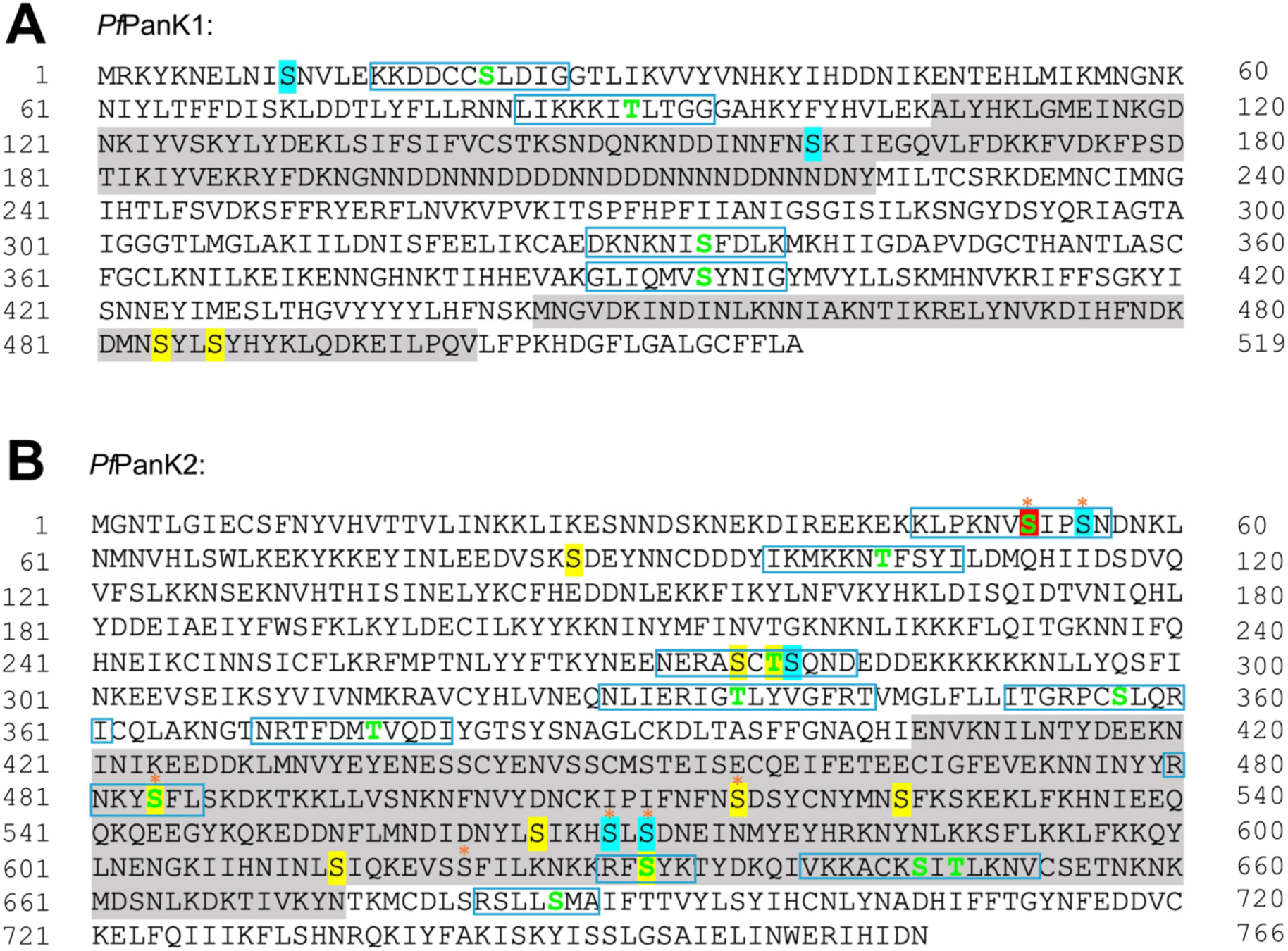
*Pf*PanK1 and *Pf*PanK2 sequences showing phosphorylated residues identified by mass spectrometry. The amino acid sequences of *Pf*PanK1 **(A)** and *Pf*PanK2 **(B)** are shown with the position of the first and last residue of each line indicated by the flanking numbers. Highlighted residues indicate the phosphorylated residues detected in the mass spectrometry analysis (yellow = detected once; cyan = detected twice; red = detected 3 times). Blue boxes indicate the 14-3-3 binding sites predicted by (50) using Eukaryotic Linear Motif (ELM), Scansite 3.0 and 14-3-3 Pred, with the phosphoserine/threonine residue required for binding indicated in bold, green. The asterisks indicate a phosphorylated residue that has been detected in one or more of the five *P. falciparum* phosphoproteomic studies. Grey-shaded regions indicate residues of *Plasmodium*-specific inserts in *Pf*PanK1 and *Pf*PanK2. Data shown are a summary of three independent analyses, each performed with a different batch of parasites.

In *Pf*PanK2, 6 out of 7 previously reported phosphorylation sites were successfully detected (S52, S55, S484, S516, S569 and S571; except S621) and an additional 8 new phosphorylation sites (S87, S276, T278, S279, S525, S565, S614 and S631) were also identified (**Figure 6B**). Out of the 14 phosphorylation sites detected in *Pf*PanK2, four were within bioinformatically-predicted *Pf*14-3-3I binding motifs, but residues predicted not to be required for binding, namely S52, T278, S484 and S631 (**Figure 6B**). Intriguingly, S55, S276 and S279, which are within the predicted *Pf*14-3-3I binding motifs, but not predicted to be required for binding, are also phosphorylated.

Phosphorylation sites within predicted *Pf*14-3-3I binding sites in *Pf*PanK2 (S52, S55, T278, S279, S484 and S631) were then mutated to investigate their role in binding to *Pf*14-3-3I. Two double mutants — S52A, S55A and T278A, S279A — were generated where multiple phosphorylation sites reside within the same predicted *Pf*14-3-3I binding site, alongside a single mutant, S484A. Although phosphorylation site S276 locates in the same predicted *Pf*14-3-3I binding site as T278 and S279, attempts to generate a mutant construct with simultaneous mutations at S276, T278, and S279 were unsuccessful. Consequently, a double mutant targeting T278 and S279 was generated instead. To increase the probability of locating the *bona fide Pf*14-3-3I binding site — particularly considering the possibility that additional phosphorylation sites may exist — we generated a *Pf*PanK2 mutant (M4-11) with 8 (T336, S357, T376, S484, S631, S646, T648 and S686) out of 11 *predicted Pf*14-3-3I binding residues mutated (excluding S52, T104 and T278). Similarly, a *Pf*PanK1 mutant (M1-4) was generated with all 4 *predicted Pf*14-3-3I binding sites (S22, T90, S334 and S394) mutated. In *Pf*PanK1-M1-4 and *Pf*PanK2-M4-11 mutants, threonine residues were mutated to valine (instead of alanine), which more closely resembles the side chain of threonine, to minimize the effect that multiple mutations may have on the protein structure. The mutated proteins were episomally expressed in *P. falciparum* with a C-terminal GFP tag as done for the WT and the earlier *Pf*PanK active site mutants.

The localisation of *Pf*PanK1 or *Pf*PanK2 and the assembly of the *Pf*PanK complex was not affected by any of the putative *Pf*14-3-3I binding site mutations introduced (**Figure S5** and **7A**). To investigate if any of the mutated sites are required for *Pf*14-3-3I binding, immunoprecipitation was performed using protein extracted from mutant *Pf*PanK1 or *Pf*PanK2-GFP parasites, with protein extracted from WT *Pf*PanK1 or *Pf*PanK2-GFP parasites serving as a control. As indicated in the anti-GFP western blots (**Figure 7B**), bands corresponding to *Pf*PanK1-GFP and *Pf*PanK2-GFP were detected in the immunoprecipitated fractions of the respective WT and mutant cell lines, providing additional evidence that all mutant proteins were successfully expressed in the transgenic parasites. When probing for 14-3-3 protein, a band (∼30 kDa) consistent with the expected size of *Pf*14-3-3I protein, was observed in the immunoprecipitated fractions prepared from the WT parasites and all mutant parasites (**Figure 7B**). For all four *Pf*PanK2 mutants, the amount of *Pf*14-3-3I protein detected in the immunoprecipitated fractions was indistinguishable from that detected in the WT samples. This suggested that none of the mutated phosphorylation sites (S52, S55, T278, S279, S484 and S631) or predicted *Pf*14-3-3I binding sites (excluding T104 which was not tested) in *Pf*PanK2 are required for *Pf*14-3-3I binding. In contrast, the M1-4 mutations in *Pf*PanK1 resulted in a substantial reduction (79 ± 5 %; mean ± SEM) in the amount of *Pf*14-3-3I detected in the immunoprecipitated samples (**Figure 7B and S6**), consistent with one (or more) of the four residues mutated being important for *Pf*14-3-3I binding. Notably, this difference in the amount of *Pf*14-3-3I bound was not due to variations in the amount of *Pf*PanK1-GFP protein in the immunoprecipitated complexes used for the analysis (**Figure S6**).

**Figure 7.**
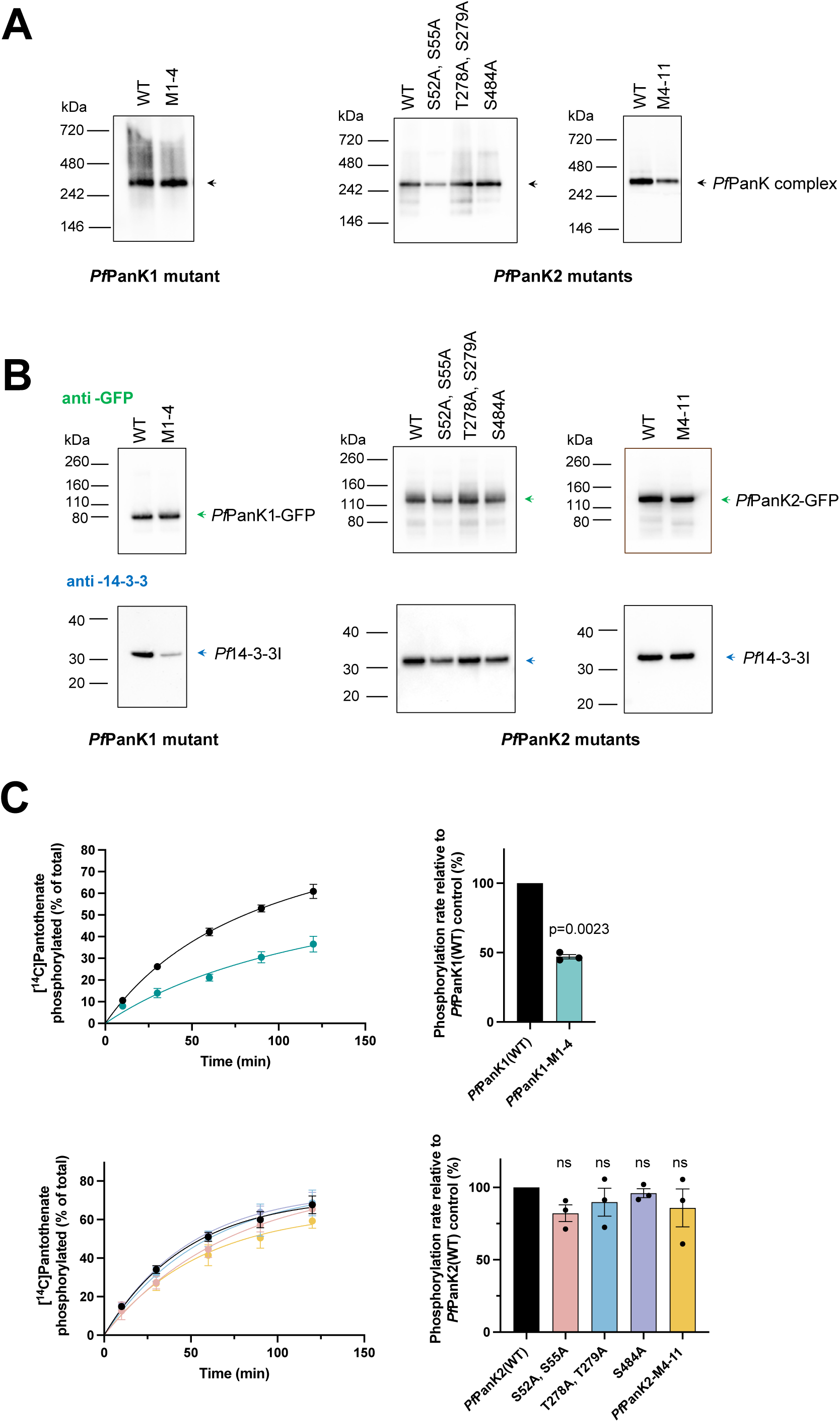
Analysis of putative *Pf*14-3-3I binding site mutant *Pf*PanK1 and *Pf*PanK2. **(A)** Anti-GFP native western blots showing the *Pf*PanK-GFP complex bands in lysates from the mutant parasites. **(B)** Anti-GFP and anti-14-3-3 denaturing western blots of the GFP-Trap immunoprecipitation fractions generated from different mutant parasites. The western blot shown is a representative of three independent experiments, each performed with a different batch of parasites. The same parasite lysate was used for panels B and C. **(C)** The phosphorylation of [^14^C]pantothenate over time (left) by GFP-Trap immunoprecipitated fractions prepared from WT and mutant *Pf*PanK1 (top), and WT and mutant *Pf*PanK2 (bottom) transgenic parasites. Phosphorylation rates (right) were determined from the overall reaction time course by one-phase association for all samples. Values are presented as a percentage relative to the WT control phosphorylation rate. Values are averaged from three independent experiments, each performed with a different batch of parasites and carried out in triplicate. Error bars represent SEM. Significant differences in phosphorylation rate are indicated with the p value (ns, not significantly different). Statistical analysis between the WT and mutant *Pf*PanK1 parasites was carried out with paired, two-tailed, Student’s t-test. Statistical analysis between the WT and mutant *Pf*PanK2 parasites was carried out with one-way analysis of variance (ANOVA) with Dunnett’s multiple comparison test. Significant differences in phosphorylation rate are indicated with the p value (ns, not significantly different).

The same immunoprecipitated samples were then tested in [^14^C]pantothenate-based phosphorylation assays to assess the PanK activity of the mutant *Pf*PanK complexes. The *Pf*PanK1-M1-4 mutant complex displayed a 53% lower (p = 0.0032, ANOVA) phosphorylation rate as compared to the WT *Pf*PanK complex (**Figure 7C**), which could be due to the significantly reduced *Pf*14-3-3I binding. However, the possibility that the reduction in PanK activity results from one (or more) of the mutations influencing the *Pf*PanK1 structure and/or stability (rather than being a result of reducing bound *Pf*14-3-3I) cannot be ruled out. The PanK activity in the immunoprecipitated samples prepared from all four *Pf*PanK2 mutants is indistinguishable (p>0.05, ANOVA) from that observed in the WT sample (**Figure 7C**), consistent with none of the residues mutated having an impact on *Pf*14-3-3I binding.

### Heterologously-expressed *Pf*PanK complex assembles *in situ* and is functional

Although we had some success at identifying the residue(s) required for *Pf*14-3-3I binding, complete elimination of *Pf*14-3-3I binding was not achieved (**Figure 7B**). Due to the conserved and multitask nature of 14-3-3 proteins, knocking out *Pf*14-3-3I is not feasible as the protein is involved in a vast number of biological processes in *P. falciparum* and is likely essential (mutagenesis index score: 0.585 and mutagenesis fitness score: -3.082) (51–54). To tackle this problem and investigate the *Pf*PanK complex in more detail, we heterologously expressed the *Pf*PanK complex in insect cells (using the baculovirus expression system (55, 56)). We utilised a FlexiBAC expression system, which offers simpler and faster recombinant protein production compared with the conventional baculovirus expression system (57).

To express all three protein components in the *Pf*PanK complex simultaneously in insect cells, all three genes of interest (GOI) were cloned into one open reading frame (ORF) and expressed under the control of the polyhedrin promoter (**Figure 8A**, **Figure S7A**). Additionally, two different 2A self-cleaving peptides (2A) were incorporated to separate the three proteins. Upon successful ribosomal skipping at the 2A junctions, three individual protein components should be obtained (**Figure 8B**). We also introduced an N-terminal His6-tag to *Pf*PanK1 and a C-terminal TwinStrep (TS) – GFP tag to *Pf*PanK2 to confirm the expression of each protein and facilitate downstream purification. Denaturing western blot analyses were performed to confirm the expression of each protein component. Protein bands corresponding to the expected sizes of His6-*Pf*PanK1, *Pf*PanK2-TS-GFP and 14-3-3 were observed, indicating all three components of the complex were co-expressed (**Figure 8C**). It was also noted that ribosomal skipping between *Pf*14-3-3I and *Pf*PanK2 by the T2A peptide was incomplete, as an additional band corresponding to the size of a fused *Pf*14-3-3I–T2A–*Pf*PanK2 protein was observed (**Figure 8C**). The incomplete skipping of 2A peptides was expected, as this has been previously documented (58, 59). Native western blot analyses were performed, and a band corresponding to the expected size for the recombinant *Pf*PanK complex was observed (slightly larger than the endogenous *P. falciparum* complex due to the presence of the additional tags) (**Figure 8D**), suggesting that the three components formed a complex *in situ*. Immunoprecipitations were then performed using GFP-Trap (*Pf*PanK2 immunoprecipitation) and nickel nitrilotriacetic acid (Ni-NTA) agarose (*Pf*PanK1 immunoprecipitation) and the activity of the immunoprecipitated proteins was assessed using a [^14^C]pantothenate-based kinase assay. Both *Pf*PanK1 and *Pf*PanK2 immunoprecipitations showed robust PanK activity, indicating that the complex had successfully assembled and was functional (**Figure 8E**).

**Figure 8.**
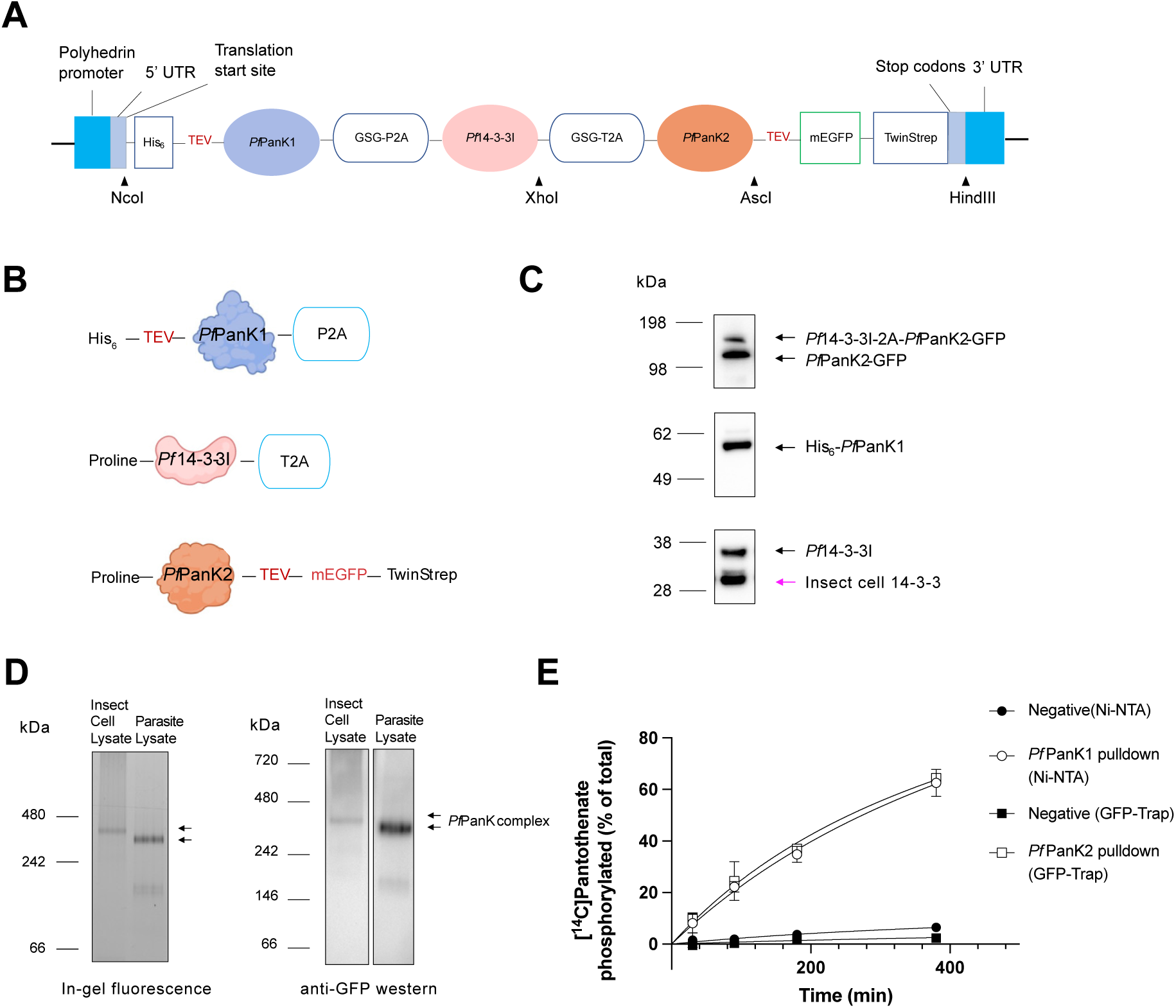
Expression of the *Pf*PanK complex in insect cells. **(A)** Overview of the expression cassette. All three GOI were cloned into the ORF *via* NcoI and HindIII restriction sites (restriction sites are shown with arrowheads). Simultaneous expression of all three proteins was achieved by incorporation of 2A peptides between genes. A glycine-serine-glycine (GSG) linker was introduced before each 2A peptide for improved skipping efficiency. Detection of target proteins and downstream purifications were facilitated by affinity tags appended to the N-terminus of *Pf*PanK1 and the C-terminus of *Pf*PanK2, respectively. A tobacco etch virus cleavage sequence was added between affinity tags and target proteins, allowing tags to be removed by TEV digestion if necessary. **(B)** Cartoon showing individual protein components expected upon successul ribosomal skipping at the 2A junctions. **(C)** Denaturing western blots using anti-GFP (top), anti-*Pf*PanK1 (middle) or pan anti-14-3-3 (bottom) antibodies showing the presence of all three components of the *Pf*PanK complex in insect cell lysates. As pan anti 14-3-3 antibodies were used, the insect cell 14-3-3 isoforms were also recognised as expected (pink arrow). **(D)** In-gel fluorescence and anti-GFP native western blots showing the fully assembled *Pf*PanK complex in insect cell lysates (left lanes) and in endogenous *P. falciparum* lysates (right lanes, detected by virtue of the C-terminal GFP tag on *Pf*PanK2). **(E)** [^14^C]Pantothenate phosphorylation by the purified *Pf*PanK complex from insect cells. Mock-purified samples from insect cells not expressing the complex were used as negative controls. Values are averaged from two independent experiments, each performed with a different batch of insect cells and carried out in triplicate. Error bars represent range/2.

In an attempt to further evaluate the expression, stability and determine the accurate molecular weight of the complexes that we heterologously expressed, FSEC was performed. In contrast to the *Pf*PanK protein complex detected in *P. falciparum* lysate (**Figure 1B**), the GFP-specific fluorescence of the heterologously expressed *Pf*PanK complex was observed in the void (retention qme around 7 min), with a tailing shoulder peak. These results indicate that majority of the protein complex we heterologously expressed is aggregated and non-functional and only a small proportion is correctly folded, which contribute to the PanK activity we previously observed in the immunoprecipitation experiments.

As recombinant proteins are expressed at very high levels under the control of the polyhedrin promoter, it is possible that overexpression of *P. falciparum* proteins resulted in aggregation of the target proteins, perhaps due to saturation of the insect cell’s chaperone systems and limited post-translational processing capabilities (60, 61). We, therefore, attempted to optimise the infection qme with virus and harvested the cells at a time point when target proteins expression just started (based on western blot analyses) to reduce the expression level. Another possible reason for the aggregation is that the composition of the extraction buffer is not optimal for the protein complex and aggregation occurs azer cell lysis. We therefore tested six stabilising additives which have been repeatedly demonstrated to increase the solubility of recombinant proteins when added during cell lysis (62). Unfortunately, even with both a reduced recombinant protein expression level and the addition of different stabilising additives, we could not prevent the aggregation from happening (**Figure S8**). Despite not being able to overcome the aggregation issue, the successful assembly and activity of a small proportion of heterologously expressed proteins prompted us to proceed with testing the importance of *Pf*14-3-3I.

### *Pf*14-3-3I is not essential for *Pf*PanK activity

To investigate the role of *Pf*14-3-3I in the *Pf*PanK complex further, we generated a recombinant virus expressing only *Pf*PanK1 and *Pf*PanK2, linked by a P2A peptide (**Figure S7B**). Denaturing western blot analyses confirmed the expression of both proteins and the absence of *Pf*14-3-3I (**Figure 9A**), however we failed to observe a complex band on native western blots (**Figure 9B**). An immunoprecipitation was performed using GFP-Trap (*Pf*PanK2 immunoprecipitation) to assess the activity of the *Pf*PanK1 + *Pf*PanK2 complex and robust PanK activity was detected in the radiolabelled assay (**Figure 9C**). We then performed a purification of the *Pf*PanK1 + *Pf*PanK2 complex using a Streptavidin affnity column targeting the TwinStrep tag. Purified fractions were tested in the radiolabelled assay and were further analysed by denaturing western (**Figure S9**). As shown in **Figure 9D**, fractions 2-4 showed a band of expected size of *Pf*PanK1, in addition to the *Pf*PanK2 band, suggesting that the complex formation between *Pf*PanK1 and *Pf*PanK2 does not require *Pf*14-3-3I. Why a band consistent with the assembly of the complex could not be detected on native western blots is unclear. It is worth noting that robust PanK activity was only detected in fractions where *Pf*PanK1 and *Pf*PanK2 both present (*i.e.* fraction 3, **Figure 9D** and **9E**). Together, these data indicate that *Pf*14-3-3I is not essential for the assembly or the function of the *Pf*PanK complex.

**Figure 9.**
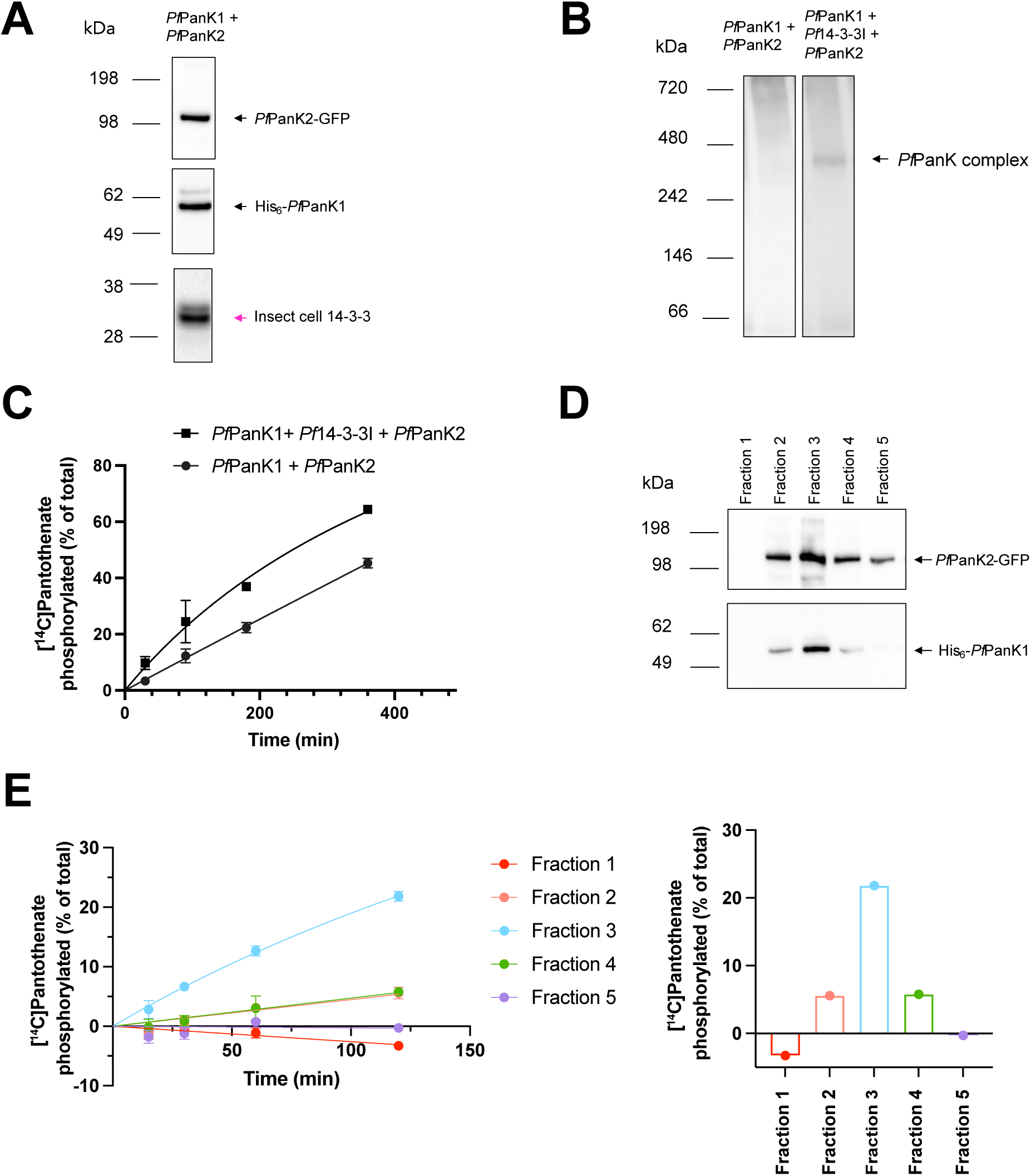
Expression of the *Pf*PanK1 + *Pf*PanK2 complex in insect cells. **(A)** Denaturing western blots using anti-GFP, anti-*Pf*PanK1 and pan anti-14-3-3 antibodies showing successful expression of *Pf*PanK2 (top) and *Pf*PanK1 (middle). Only the insect cell 14-3-3 isoforms are detected (bottom, pink arrow) **(B)** Anti-GFP native western blots showing the attempt to detect the *Pf*PanK1 + *Pf*PanK2 complex (left) and the *Pf*PanK complex (with *Pf*14-3-3I included, right) in insect cell lysates. **(C)** [^14^C]Pantothenate phosphorylation by the purified *Pf*PanK complex (including *Pf*14-3-3I) and *Pf*PanK1 + *Pf*PanK2 (excluding *Pf*14-3-3I) from insect cells. Values are averaged from two independent experiments, each performed with a different batch of insect cells and carried out in triplicate. Error bars represent range/2. **(D)** Denaturing western blots using anti-GFP (top) or anti-*Pf*PanK1 antibody (bottom) showing the presence of both *Pf*PanK2 (top) and *Pf*PanK1 (bottom) in Strep-purified (*Pf*PanK2-purified) protein fractions 2-4, in which PanK activity was detected. **(E)** The phosphorylation of [^14^C]pantothenate over time (left) by each purified fraction in **(D)**. The percertage of [^14^C]pantothenate phosphorylated at the end of 120 min (right) by each purified fraction. These data are representative of two independent experiments. Error bars represent range/2.

## Discussion

PanKs plays a pivotal role in CoA biosynthesis and usually function as a homodimers (24–31). Recently, apicomplexan parasites were shown to harbour heteromeric PanK complexes, (33, 63). Based on the comparison of amino acid sequences it was proposed that PanKs from *Plasmodium* and *Toxoplasma* parasites are unlikely to function as homodimers and only the heteromeric PanK1/PanK2 complex can serve as a functional PanK (33).

Using advanced AI-based structure-prediction algorithms, we were able to predict the structure of the *Pf*PanK molecular ensemble in complex with its ligands. Further docking analysis supported the hypothesis that only one of the two putative active sites in the *Pf*PanK complex is able to perform the kinase reaction (*i.e.* the complete active site; **Figure 2D**). Results from our site-directed mutagenesis experiments were consistent with this hypothesis. Mutation of critical residues in the complete active site resulted in elimination or a significant reduction in PanK activity whereas mutations of residues (apart from I343) in the incomplete active site had no effect on PanK activity (**Figure 3B** and **3C**). Although situated in the incomplete site, mutation of I343 to valine in *Pf*PanK1 resulted in a 40% reduction of PanK activity. It is unlikely that the I343V mutation directly influences the stability of the complete active site of the *Pf*PanK complex or interferes with the catalysis of pantothenate given mutation of none of the other residues in the site had an effect. It is possible that the mutation results in a local structure change (potentially due to change in bulkiness), which could influence the stability of the *Pf*PanK complex, resulting in a reduced phosphorylation rate. In addition to stabilising the active site, E233 of *Pf*PanK1 is the predicted catalytic base essential for phosphate transfer in the kinase reaction, and it also contributes to the binding of magnesium (42). Our results with the E233V mutant are consistent with work showing that mutating the equivalent residue in *Hs*PanK3 (E138) to valine abolishes *Hs*PanK3 activity (64). The consequences of the R207W substitution of *Hs*PanK3 are less catastrophic but result in decreased binding affinity for both pantothenate and the allosteric inhibitor, acetyl-CoA (64). Mutating the equivalent residue (R295W) in *Pf*PanK1 resulted in reduced enzyme activity, but the enzyme was also still active. Overall, our results are consistent with the *P. falciparum* PanK complex having only one functional active site composed of residues from both *Pf*PanK1 and *Pf*PanK2. This demonstrates, for the first time, that *Pf*PanK2 contributes to the parasite PanK activity.

*Pf*PanK1 is important for normal intraerythrocytic proliferation of *P.falciparum* (20). In this study we show, using knockdown experiments, that *Pf*PanK2 is also essential for normal intraerythrocytic proliferation of *P. falciparum* (**Figure 5B** and **5C**). This is consistent with both *Pf*PanKs being required for the formation of a functional *Pf*PanK complex. Our results are seemingly at odds with findings that both PanKs are dispensable during the intraerythrocytic stage of *Plasmodium berghei* and *Plasmodium yoelii* (65–67). This could be explained by the fact that these parasites preferentially infect reticulocytes (68, 69), which have been shown to provide a rich pool of nutrients for the parasites (66). It is therefore possible that the reticulocyte-residing parasites can salvage sufficient CoA or CoA intermediates from the host cell, whereas *P. falciparum* residing within more mature erythrocytes cannot, making the two *Pf*PanKs indispensable, as proposed by Srivastava *et al.* (66) and Tjhin *et al.* (20).

From the phosphoproteomic experiments, we identified four new phosphorylation sites in *Pf*PanK1 and an additional eight new phosphorylation sites in *Pf*PanK2 (**Figure 6**). Out of all *Pf*PanK phosphorylation sites detected in this study and in previous publications (45–49), none are within the *Pf*14-3-3I binding sites predicted for *Pf*PanK1. By contrast, seven phosphorylation sites reside within the predicted *Pf*14-3-3I binding sites in *Pf*PanK2, of which four are predicted to be important for binding. Mutagenesis data showed that 10 out of 11 (except T104 which was not tested) predicted *Pf*14-3-3I binding residues in *Pf*PanK2 are not required for *Pf*14-3-3I binding (**Figure 7B**). On the other hand, mutation of four predicted 14-3-3 binding sites (S22, T90, S334 and S394) in *Pf*PanK1 resulted in a significant reduction in the amount of *Pf*14-3-3I bound to the *Pf*PanK complex (**Figure 7B**). Using the AlphaFold-predicted *Pf*PanK1-*Pf*PanK2 structure (**Figure 2C**), we envisaged that S334 is the most likely *Pf*14-3-3I binding residue on the basis that (i) S22 and T90 are situated in close vicinity to the functional active site and the phosphate-binding loop (P-loop, which plays a vital role in the binding and positioning ATP for catalysis) of the complex and binding of *Pf*14-3-3I in this region would likely interfere and/or impair the kinase reaction (**Figure S9**), and (ii) S394 is located in the ⍺ helix that sits at the predicted dimer interface and is positioned close to the N396, M400, L403 and M407 residues (all located in the ⍺-helix), which were found to be important for *Hs*PanK3 dimerisation (42). Therefore, binding of *Pf*14-3-3I to S394 might disrupt the dimerisation of *Pf*PanK1 and *Pf*PanK2. Whether S334 is a *bona fide Pf*14-3-3I binding site remains to be determined.

Although many known 14-3-3 binding proteins contain a single binding site, approximately 50 proteins have more than one confirmed 14-3-3 binding sites (70–73). The fact that we did not observe complete elimination of *Pf*14-3-3I binding in the *Pf*PanK1-M1-4 mutant raises the possibility that there is more than one *Pf*14-3-3I binding site on the *Pf*PanK complex. Although 14-3-3 monomers are less stable than dimers, they can retain some functions of the dimeric 14-3-3 and may harbour additional functional roles due to their exposed inter-subunit surfaces (41, 74). It is possible that a *Pf*14-3-3I monomer (instead of a dimer) might still be able to bind to other binding site(s) in the *Pf*PanK1-M1-4 mutant and the change in the complex size due to loss of one *Pf*14-3-3I monomer (∼30 kDa) might not be resolved on NativePage (**Figure 7A**).

A known function of 14-3-3 proteins is to regulate the intracellular localisation of their binding partners by interfering with the function of a nearby targeting sequence (38, 75, 76). Our *Pf*PanK1-M1-4 mutant largely ruled out the possibility that *Pf*14-3-3I regulates the cellular localisation of the *Pf*PanK complex (**Figure S5**). It is also unlikely that *Pf*14-3-3I acts as a bridge between *Pf*PanK1 and *Pf*PanK2 since (i) there is no change in the *Pf*PanK complex size observed in the *Pf*PanK1-M1-4 mutant (**Figure 7A**), and (ii) the heterologously expressed *Pf*PanK1 and *Pf*PanK2 assembled without *Pf*14-3-3I, although we cannot rule out the possibility that the insect cell 14-3-3 is used for assembly (**Figure 9A**). Additionally, the fact that the heterologously expressed *Pf*PanK1-*Pf*PanK2 assembly is catalytically active points to *Pf*14-3-3I being non-essential for *Pf*PanK activity (**Figure 9C**). It is therefore likely that *Pf*14-3-3I plays a role in maintaining the stability of the *Pf*PanK complex, as has been previously reported for other proteins (77–79). This might also explain why we were unable to detect a complex band for the heterologously expressed *Pf*PanK1-*Pf*PanK2 assembly. Despite being catalytically active, it is possible the *Pf*PanK1-*Pf*PanK2 assembly is less stable without the presence of *Pf*14-3-3I. If this is the case, a possibility for why we did not detect a change in complex size in the *Pf*PanK1-M1-4 mutant might be that the *Pf*PanK1-*Pf*PanK2 assembly is intrinsically unstable and cannot be detected under native conditions (similar to what has been observed with the *Pf*PanK1-*Pf*PanK2 assembly in insect cells). Therefore, only the *Pf*PanK complex with the presence of *Pf*14-3-3I was detected in **Figure 7A**, whereas the *Pf*PanK1-*Pf*PanK2 assembly (without *Pf*14-3-3I) was not detected.

Our inset cell data suggest that the *Pf*PanK complex could be heterologously expressed and assembled, however, the majority of the recombinant proteins expressed are aggregated and non-functional, with only a small proportion being correctly assembled and functional (**Figure S8**). Although we have thus far been unable to overcome the aggregation issue, it is the first time we have been able to heterologously express a functional *Pf*PanK complex. This breakthrough has allowed us to study the *Pf*PanK complex without *Pf*14-3-3I and determine that *Pf*14-3-3I is not required for the complex to be functional, which could not be achieved endogenously. It is unlikely that the insect cell 14-3-3 was used for assembly in the absence of *Pf*14-3-3I as we did not observe the *Pf*PanK complex band in **Figure 9B**. It is possible that subtle structural differences between *P. falciparum* and insect cell 14-3-3 proteins may impair proper interaction with *Pf*PanK subunits, or that post-translational modifications present in *P. falciparum* but absent in insect cells are required for correct complex assembly. Our *P. falciparum* mutagenesis data clearly indicate that both *Pf*PanK1 and *Pf*PanK2 are required for the function of the *Pf*PanK complex, with residues from both *Pf*PanK1 and *Pf*PanK2 contributing to the stabilisation of the active site and therefore, the parasite’s PanK activity.

## Methods

### Malaria parasite culture

Culturing of intraerythrocytic stage *P. falciparum* parasites (3D7 strain and transgenic cell lines generated in this study, which were derived from 3D7) was carried out as previously described (80, 81). Parasites were maintained within human erythrocytes (typically Group O^+^, Rh^+^, 4% haematocrit) in RPMI-1640 medium (Thermo Fisher Scientific, catalogue number 72400120) supplemented with 25 mM HEPES, 11 mM glucose, 200 μM hypoxanthine, 24 μg/mL gentamicin and 6 g/L Albumax II on a horizontally rotating shaking incubator (Ratek) at 37°C under an atmosphere of 96% nitrogen, 3% carbon dioxide and 1% oxygen.

### Construction of *P. falciparum* plasmids

To amplify the mutant *PfpanK1* and *PfpanK2* DNA fragments, a MegaPrimer PCR approach was used (82, 83) (**Table S1**), with *Pf*PanK1-pGlux-1 or *Pf*PanK2-pGlux-1 plasmid serving as template. To generate *Pf*PanK2-M4-11-pGlux-1, *Pf*PanK2-S484A-pGlux-1 plasmid was used as the template. PCR reactions were performed using PrimeSTAR GXL DNA Polymerase (Takara). Purified PCR products and pGlux-1 plasmid DNA (provided by Prof Alexander Maier, Australian National University (84)) were digested using XhoI and KpnI restriction endonucleases and subsequently assembled using an In-Fusion HD Cloning Kit (Takara). Ligated constructs, conferring ampicillin resistance, were transformed into Stellar chemically-competent cells (Takara), miniprepped and verified by Sanger sequencing (**Table S1**).

To generate the *Pf*PanK1 or *Pf*PanK2-GFP-KD construct, a C-terminal homologous region (HR, 992 base pairs of the *PfpanK1* gene or 1,165 base pairs of the *PfpanK2* gene) was amplified from the *Pf*PanK1 or *Pf*PanK2-pGlux-1 plasmid DNA with restriction endonuclease sites, SpeI and KpnI, incorporated (**Table S1**). The SpeI/KpnI-digested PCR product, was then ligated into digested eGFP-*glmS* plasmid (provided by Parichat Prommana, National Center for Genetic Engineering and Biotechnology (85)). Ligated constructs, conferring ampicillin resistance, were transformed into RbCl-competent DH5α *E. coli*, miniprepped and verified by Sanger sequencing (**Table S1**).

### Parasite transfection

All constructs used for transfection were maxiprepped using a Qiagen HiSpeed Plasmid Maxi Kit. A 100 µg aliquot of the desired construct was transfected into young, ring-stage *P. falciparum* as previously described (84, 86). Electroporation was carried out using a BioRad Gene Pulser and Pulse Controller Unit (resistance at ∞ Ohms, 310 kV, high capacitance, 950 µF). Electroporated cells were then transferred to culture flasks containing prewarmed medium and uninfected erythrocyte (haematocrit 3%). Six hours after transfection, WR99210 (10 nM) was added to the cultures to begin positive selection for mutant *Pf*PanK1-pGlux-1 or *Pf*PanK2-pGlux-1 transfected parasites. Stable transgenic populations were obtained three weeks post-transfection and were maintained in the presence of WR99210 to prevent the loss of the mutant plasmids.

### Generation of 3D7-*Pf*PanK1/2-GFP-*glmS* and CJ-A-*Pf*PanK2-GFP-*glmS* parasites

A conventional drug cycling approach was used to generate the 3D7-*Pf*PanK1/2-GFP-*glmS* (for endogenous expression of *Pf*PanK1-GFP and *Pf*PanK2-GFP) and CJ-A-*Pf*PanK2-GFP-*glmS* parasites (for assessing the essentiality of *Pf*PanK2). After transfection with the *Pf*PanK1 or *Pf*PanK2-GFP-KD construct, blasticidin S HCl (2 μg/mL) was first added to select for parasites carrying the construct episomally. Blasticidin-resistant 3D7 or CJ-A transfectant parasites were then maintained in the absence of blasticidin S HCl for two weeks to promote the loss of episomal copies of the *Pf*PanK1 or *Pf*PanK2-GFP-KD construct. Following this, to select for parasites that retained the blasticidin selection cassette *via* integration into the genome, blasticidin S HCl (2 μg/mL) was re-introduced to parasite cultures. Subsequently, the two-week on/off cycling was performed another two times. A clonal population of each integrant parasite was isolated by serial dilution cloning as previously described (87). Genomic DNA from cloned parasites was extracted from isolated trophozoite stage *P. falciparum* parasites using a DNeasy Plant Mini Kit (QIAGEN) following the manufacturer’s instructions. Integration of the *Pf*PanK1 or *Pf*PanK2-GFP-KD construct and the clonality of the parasites were assessed *via* diagnostic PCR (**Table S1**).

### Insect cell culture

Sf9 insect cells were maintained at 27 °C in suspension with shaking at 140 rpm in Insect-XPRESS medium (Lonza, catalogue number 12-730Q) supplemented with 100 U/mL penicillin and 0.1 mg/mL streptomycin. Cells were passaged twice a week and maintained at ≥95% cell viability (quantified by trypan blue staining).

### Construction of recombinant baculovirus

To generate the *Pf*PanK-pOCC shuttle vector, the pOCC5 shuttle vector (Plasmid #118863, Addgene) was first modified to incorporate a C-terminal mEGFP-TwinStrep cassette using restriction endonuclease sites AscI and HindIII. The desired *Pf*PanK ORF sequence (**Figure 8A**) was codon-optimised, synthesised and subsequently cloned into the pOCC5-mEGFP-TwinStrep plasmid (between the NcoI and AscI sites) by Genscript to generate the *Pf*PanK-pOCC construct (**Table S1 and S2**, **Figure S7A**). To then generate the bicistronic vector that only expresses *Pf*PanK1 and *Pf*PanK2 (*i.e.* no *Pf*14-3-3I), the His_6_-*Pf*PanK1 ORF was amplified with restriction endonuclease sites NcoI and Xhol incorporated (**Table S1**). Purified PCR products and *Pf*PanK-pOCC plasmid DNA were digested using NcoI and XhoI restriction endonucleases. Ligated constructs, conferring ampicillin resistance, were transformed into competent Top10 *E. coli*, miniprepped and verified by Sanger sequencing (**Table S1**, **Figure S7B**).

DefBac bacmid (provided by Régis P Lemaitre, Max Planck Institute of Molecular Cell Biology and Genetics) was transformed into competent DH10β *E. coli* and maxiprepped using a Nucleobond Xtra Maxi kit (Scientifix). Recombinant viruses were generated as detailed by Lemaitre *et al.* (57). P1 and P2 virus stocks were filtered through a 0.22 µm filter and stored at 4°C protected from light.

### Baculovirus expression of *Pf*PanK proteins in Sf9 cells

For expression of the *Pf*PanK complex or the *Pf*PanK1-*Pf*PanK2 assembly, exponentially growing Sf9 cells (3 × 10^6^ cells/mL) were infected by the direct addition of P2 (0.2% v/v) virus stock. Azer the virus was added, the cells were allowed to stand for 1 h at 27°C without shaking. After 1 h, the virus + cell suspension were diluted to a final density of 1.5 × 10^6^ cells/mL. The cells were incubated for 4-5 days shaking at 27°C. Before harvesting, the cells were checked for infection by staining with trypan blue (∼50% viability). Infected Sf9 cells were centrifuged at 4,500 × g for 5 min and the cell pellets were snap frozen with liquid nitrogen and stored at -80°C until further use.

### Streptavidin purification

Infected Sf9 cell pellets were resuspended in binding buffer (100 mM Tris-HCl, 150 mM NaCl, 1 mM EDTA, pH 8) supplemented with DNase and protease inhibitor cocktail (PIC, Sigma-Aldrich) and were subsequently mechanically lysed with an EmulsiFlex-C5 high pressure homogeniser (AVESTIN). Cells were passed through the pressure cell two times in total at a pressure of ∼10,000 psi to ensure complete lysis and immediately centrifuged (30,000 × g, 30 min, 4°C) to pellet cell debris and insoluble proteins. The clarified lysate was then filtered through 0.2 µm filters and loaded onto a 5 mL StrepTrap^TM^ HP column (Cytiva) pre-equilibrated with binding buffer at a 1 mL/min flow rate. The column was then washed with 10 column volumes of binding buffer. Proteins were eluted in a linear gradient of buffer A (100 mM Tris-HCl, 150 mM NaCl, 1 mM EDTA, 2.5 mM desthiobiotin, pH 8) to buffer B (100 mM Tris-HCl, 150 mM NaCl, 1 mM EDTA, 7.5 mM desthiobiotin, pH 8) over 6 CV at a 1 mL/min flow rate. Purified samples were then analysed by SDS-PAGE and western blots.

### Fluorescence microscopy

*P. falciparum*-infected erythrocytes expressing WT or mutant *Pf*PanK1-GFP or *Pf*PanK2-GFP fusion proteins were observed with an Applied Precision DeltaVision microscope. Trophozoite or schizont stage *P. falciparum*-infected erythrocytes were pelleted (2,000 × g, 30 s) and resuspended in malaria saline (125 mM NaCl, 5 mM KCl, 25 mM HEPES, 20 mM glucose and 1 mM MgCl_2_, pH 7.4) containing 2 μg/mL Hoechst 33258. Cells were incubated in the dye for 15 min at 37°C, subsequently pelleted and washed two times with malaria saline prior to imaging. Images were captured using a 100× oil-immersion objective, deconvoluted and linearly adjusted for contrast and brightness.

### Parasite growth assay

The parasite growth assays were performed using a SYBR Safe-based fluorescence assay as previously described (14, 88). Synchronous ring-stage CJ-A parent parasites and CJ-A-*Pf*PanK2-KD parasites were seeded in triplicate at 0.5% parasitaemia and 1% haematocrit. Parasites were incubated for 96 h in the presence of various concentrations of GlcN. For the *in vitro* pantothenate requirement experiments and the low-pantothenate experiments, cells were washed twice in complete pantothenate-free medium (1,500 × g, 5 min) to remove any pantothenate before seàng up experiments.

To monitor parasite growth over multiple cycles, FACS analysis was performed with *P. falciparum*-infected erythrocytes using a BD LSR II flow cytometer. Live cells were stained with Hoechst 33258 nuclear stain as described previously and were then diluted in malaria saline to a concentration of ∼10^6^ – 10^7^ cells/mL. One hundred thousand cells were sampled from each tube at a low sampling speed with the following settings: forward scatter = 308 V (log scale), side scatter = 308 V (log scale), Alexa Fluor 488 = 559 V (log scale) and Pacific Blue = 478 V (log scale). An uninfected erythrocyte sample was first analysed to establish a gating strategy that defined a threshold above which erythrocytes were deemed to be infected with *P. falciparum*.

### Denaturing western blots

To check for the expression of all *Pf*PanK components in insect cells, virally-infected insect cells were lysed in 1% (w/v) sodium dodecyl sulfate (SDS), passed through a 25-gauge needle (20×) and centrifuged (16,000 × g, 30 min, 4°C) to pellet insoluble materials. The supernatant containing the soluble protein was then supplemented with 1× NuPAGE^TM^ LDS Sample Buffer and 1× NuPAGE^TM^ Sample Reducing Agent, vortexed to mix and then heated at 95°C for 10 min. To confirm the expression of endogenously-tagged and mutant *Pf*PanK1 and *Pf*PanK2 in *P. falciparum*, trophozoite-stage parasites were saponin-isolated (89) and lysed as previously described (33). Protein samples generated from both *P. falciparum* parasites and insect cells were separated on NuPAGE^TM^ Novex 4–12% Bis-Tris gels (Invitrogen) and transferred onto nitrocellulose membranes as previously described (33). The membrane was blocked with blocking buffer (4% (w/v) skim milk in tris-buffered saline with 0.1% v/v Tween 20 (TBST)) overnight at 4°C with constant shaking.

For immunodetection of the desired proteins, blocked membranes were exposed to specific primary and secondary antibodies (diluted in blocking buffer) for 1.5 h at room temperature (RT). The following primary antibodies were used: mouse anti-GFP monoclonal antibody (0.4 μg/mL final concentration; Roche; REF 11814460001), rabbit pan-specific anti-14-3-3 polyclonal antibody (0.5 μg/mL final concentration; Abcam14112), mouse anti-heat shock protein 70 (HSP70) monoclonal antibody (0.5 μg/mL final concentration; provided by Prof Alexander Maier) and rabbit anti-*Pf*PanK1 final bleed (1:5000 dilution, provided by Patrick A M Jansen, Radboud University Medical Center (15)). The secondary antibodies used were an anti-mouse goat horseradish peroxidase (HRP)-conjugated antibody (1:5000 dilution; Abcam; ab6789) or an anti-rabbit goat HRP-conjugated antibody (1:2500 dilution; Thermo Fisher Scientific, A16096). To visualise the protein band(s), membranes were exposed to the Pierce ECL Western Blotting Substrate (Thermo Fisher Scientific). After a 5 min incubation with the substrate, membranes were visualised with a ChemiDoc^TM^ MP Imaging System (Bio-Rad).

### Native western blots

Parasite lysates were prepared from saponin-isolated mature trophozoite stage parasites as described by Tjhin *et al.* (33), with modifications. Briefly, washed saponin-isolated parasites were suspended in 500 µL of native lysis buffer (10 mM Tris-HCl, pH 7.4, 140 mM NaCl, 10 × PIC, 0.5% (w/v) digitonin, 60 units of benzonase nuclease and 4 mM MgCl_2_). The resuspended cells were incubated at 4°C with constant mixing for 30 min and subsequently centrifuged at 16,000 × g for 30 min at 4°C to remove haemozoin and other cell debris. Protein samples were diluted in 1× NativePAGE^TM^ Sample Buffer and supplemented with 10% (v/v) loading dye (50% glycerol, 0.1% Ponceau S). Insect cell lysates were prepared in the same way. Clear native electrophoresis was performed using NativePAGE^TM^ (4–16%) gels (ThermoFisher). Immunodetection was performed in the same manner as described for denaturing western blots.

### Immunoprecipitations

To purify GFP-tagged proteins from parasite or insect cell lysates, immunoprecipitation was performed using GFP-Trap Agarose (Proteintech). Saponin-isolated trophozoite-stage parasites or insect cells were resuspended in 500 μL of cold GFP-Trap lysis buffer (10 mM Tris-HCl, pH 7.5, 150 mM NaCl, 0.5 mM EDTA, 10 × PIC, 0.5% (w/v) digitonin, 60 units of benzonase nuclease and 4 mM MgCl_2_). Cells were lysed and haemozoin and other cell debris removed as described in the previous section.

To purify His-tagged proteins from insect cell lysates, immunoprecipitation was performed using Ni-NTA Agarose (QIAGEN). Insect cells were resuspended in 500 μL of cold Ni-NTA lysis buffer (50 mM Tris-HCl, 500 mM NaCl, 10 mM imidazole, 10 % glycerol, pH 7.5), 0.5 % (w/v) digitonin, 10× PIC, 60 units of benzonase nuclease and 4 mM MgCl_2_). Cells were lysed at 4°C with constant mixing for 30 min and subsequently centrifuged at 16,000 × g for 30 min at 4°C to remove cell debris and insoluble proteins. The supernatant was then transferred to a new tube. The Ni-NTA slurry (25 μL) was first equilibrated with 20 volumes of cold Ni-NTA binding buffer, the bead suspension centrifuged (2,500 × g, 2 min, 4°C) and the supernatant removed. The wash was performed another two times before the cell lysate was added to the Ni-NTA beads and incubated for 1 h at 4°C with constant mixing. Subsequently, the beads were washed four times with cold wash buffer as described previously in the equilibration step, to remove any non-specific, low affinity binding proteins, prior to downstream applications.

### [^14^C]Pantothenate phosphorylation assay

Phosphorylation of [^14^C]pantothenate was determined using a combination of zinc sulphate (ZnSO_4_) and barium hydroxide (Ba(OH)_2_) (Somogyi reagent), which precipitates phosphorylated compounds from solution, as previously described (33, 90). Each reaction contained a final concentration of 50 mM Tris-HCl (pH 7.4), 5 mM ATP, 5 mM MgCl_2_ and 2 μM (0.1 μCi/mL) [^14^C]pantothenate. Reaction mixtures were incubated at 37°C throughout the experiment and reactions were initiated by the addition of the immunoprecipitated samples from *P. falciparum* or insect cells. At pre-determined time points, 50 μL of the reaction mixture was transferred to wells of a 96-well filter plate, each with a 0.2 μm hydrophilic polyvinylidene fluoride (PVDF) membrane filter (Corning), which had been pre-loaded with 50 μL 150 mM Ba(OH)_2_. Phosphorylated compounds in each well were precipitated by the addition of 50 μL 150 mM ZnSO_4_ to generate the Somogyi reagent (91), the wells processed, and the radioactivity in the plate determined as detailed previously (90). To determine the total radioactivity in each phosphorylation assay, 50 μL of each reaction was added in the wells of an OptiPlate-96 microplate (PerkinElmer), thoroughly mixed (by pipeàng at least 50 times) with 150 μL of Microscint-40 (PerkinElmer) and measured as detailed previously (90).

### Fluorescence-coupled size exclusion chromatography

Parasite and insect cell lysates were prepared for FSEC analysis as described above for native western blots. Lysates were filtered through a 0.22 µm filter prior to loading onto a Superdex 200 5/150 GL column (GE healthcare) attached to a Dionex UltiMate^TM^ 3000 UHPLC (Thermo Scientific) coupled to a fluorescence detector. To generate the molecular weight calibration curve, a gel filtration markers kit (MWGF1000, Sigma-Aldrich) was used. The absorbance of each protein standard was monitored at 280 nm. Fluorescence of the GFP-tagged proteins was monitored with an excitation wavelength at 488 nm and an emission wavelength at 509 nm. A 10 mM Tris-HCl, 140 mM NaCl, pH 7.40 buffer was used, and the chromatography was performed at a 0.15 mL/min flow rate for 30 min at RT. For trouble-shooting aggregation in insect cells, a 50 mM Tris-HCl, 500 mM NaCl, 10 % glycerol, pH 7.5 buffer was used.

### Phospho-peptide enrichment and mass spectrometry

To identify *Pf*PanK phosphorylation sites *via* mass spectrometry, *Pf*PanK-GFP complex was immunoprecipitated from *Pf*PanK1-GFP parasites (1 L of *P. falciparum* culture, 4% haematocrit, 5 % parasitemia) using the ChromoTek GFP-Trap Magnetic Agarose (Proteintech) in the presence of 2 × PhosSTOP phosphatase inhibitor (Sigma-Aldrich, catalogue number 4906845001). After immunoprecipitation, the bound fractions were magnetically pelleted, snap frozen with liquid nitrogen and sent (on dry ice) to the Australian Proteomics Analysis Facility (NSW) for processing and mass spectrometry analysis.

The magnetic beads with the protein complexes bound were reconstituted in 100 mM triethylammonium bicarbonate (TEAB) buffer. Proteins were reduced with DTT (10 mM) at 60 °C for 45 min with rigorous shaking followed by alkylation with indole-3-acetic acid (IAA, 25 mM) in the dark at room temperature for 45 min with rigorous shaking. Proteins were digested with 1 μg of trypsin overnight with rigorous shaking. Beads were centrifuged at 14,100 × g for 5 min, followed by the exposure to a magnetic rack to separate the beads from the solution. Lastly, the supernatant solution containing peptides was transferred to a new tube. About one tenth of the peptide solution was retained for MS analysis before performing any further enrichment.

For the phospho-peptide enrichment procedure, approximately 50 μg TiO_2_ bead slurry was packed into a pipette tip prepacked with glass fibres. The TiO_2_ beads were activated using GTA binding buffer (80% acetonitrile (ACN), 5% trifluoroacetic acid (TFA), 80 mg/mL glycolic acid). Simultaneously, proteolytic peptides from the sample were diluted with GTA binding buffer. Following acidification, peptides were loaded onto the TiO_2_ bead packed tip and allowed to bind to the beads. The unbound peptides (flow-through fraction) were collected and retained for MS analysis. Following binding, two sequential washes were performed to remove non-specifically bound peptides, first with washing buffer-1 (80% ACN, 1% TFA) twice, followed by washing buffer-2 (10% ACN, 0.1% TFA) twice. Finally, the bound phospho-peptides were eluted using ammonia solution and acidified with formic acid for mass spectrometry analysis. Samples were stored at -20 °C until use. Prior to LC-MS/MS analysis, samples were centrifuged at 14,100 × g for 5 min and half of the supernatant sample was subjected to 1D-IDA nano-LC MS/MS analysis (IDA-LC–MS/MS).

Peptide samples were injected onto the peptide trap column (C18 PepMap 100, 100 Å, 5 μm, 300 μm × 5 mm) and desalted with loading buffer (99.9% water, 0.1% formic acid). Peptides were eluted from the trap into an in-house packed analytical column (Halo C18, 160 Å, 2.7 μm, 100 μm × 30 cm) with the linear gradients of mobile phase A (99.9% water, 0.1% formic acid) and B (99.9% acetonitrile, 0.1% formic acid): mobile phase B (25%) over 85 min with a flow rate of 600 nL/min across the gradient. The eluent from the trap was separated over the analytical column. The column eluent was directed into the ionization source of the mass spectrometer (Q-Exactive HFX, Thermo Fisher Scientific). A 2.6 kV electrospray voltage was applied *via* a liquid junction upstream of the column. Peptide precursors from 350 to 1850 m/z were scanned at 60 k resolution with an AGC target value of 3 × 10^6^. The 20 most intense ions from the preceding survey scan were fragmented by Higher-Energy Collisional Dissociation (HCD) using normalized collision energy of 28 with an isolation width of 1.4 m/z. Only precursors with charge state +2 to +5 were selected for MS/MS analysis. The MS method had a minimum signal required value of 3×10^3^ for MS2 triggering, an automatic gain control (AGC) target value of 1 × 10^5^ for MS2 and a maximum injection qme of 70 ms for MS2. MS/MS scan resolution was set at 15 k. The dynamic exclusion was set to 30 s.

The mass spectrometric data for each sample set were searched using Proteome Discoverer (version 2.1, Thermo Scientific). The data were processed using search engines SequestHT and Mascot (Matrix Science, London, UK) against all *P. falciparum* (3D7) protein sequences downloaded from the Uniprot database. The search parameters for the data processing were as follows: Enzyme Name: Trypsin, Maximum missed cleavages: 2, Precursor mass tolerance: 10 ppm. Fragment mass tolerance: 0.02 Da, Dynamic modifications: Oxidation (M), Deamidated (N, Q), PyroGlu (Q), Acetyl (Protein N-Terminus) and Phospho (STY). Static Modification: Carbamidomethyl (C). Phospho-site localization probability was calculated using an inbuilt feature of Proteome Discoverer 2.1; ptmRS. False discovery rate (FDR) and result display filters were set as follows: Protein (SEQUEST), Peptide (search engine specific) and PSM FDR<1%, Master proteins only.

### Structural prediction of the *Pf*PanK heteromer and molecular docking

AlphaFold 2 was used to predict the *Pf*PanK1-*Pf*PanK2 heteromer structure. The full-length sequences of *Pf*PanK1 (Q8ILP4) and *Pf*PanK2 (Q8IL92) were used. Five structures were predicted for the protein complex, which were subsequently assessed using pLDDTs scores. The top-ranked predicted structure was then analysed using PyMOL 2.5.4.

The 3D structure of each substrate was prepared using the Ligprep module in Maestro (Schrödinger, LCC, New York, NY) with the OPLS_3e force field (92). All possible stereoisomers and relevant protonation states were determined using the Epik module. The heteromer structure was prepared using the Protein Preparation Wizard module (Schrödinger, LCC, New York, NY). All hydrogen atoms were added, and bond orders were assigned. All structures were refined by running a restrained minimisation to optimise the hydrogen bond network with the OPLS_3e force field. Molecular docking was performed using Glide in Schrödinger. For the complete active site, the coordinates of substrates in the homodimeric *Hs*PanK3 structure (PDB code: 5KPR) were defined as the center of the grid of the *Pf*PanK1-*Pf*PanK2 complex. For the incomplete active site, the position of the key residues was defined as the center of the grid. The outer box size was set to 20 Å, and the inner box size was 10 Å, to accommodate the residues of the active site. Standard precision (SP) was used for docking and the scoring function GlideScore was applied for ranking poses. For each docking simulation, 10 poses were written out followed by post-docking minimisation, and the pose with the highest score was used for subsequent analysis.

### Statistical analysis

To test for statistically significant differences between the means of pantothenate phosphorylation rates of more than two groups, one-way analysis of variance (ANOVA) with Dunnett’s multiple comparison test was performed using GraphPad Prism 10. Statistical analysis between the CJ-A parent parasites and the CJ-A-*Pf*PanK2-*glmS* parasites was carried out with unpaired, two-tailed, Student’s t-test using GraphPad Prism 10.

## Acknowledgements

We would like to thank the Canberra Branch of the Australian Red Cross Lifeblood for the provision of red blood cells. We are grateful to the Australian Proteome Analysis Facility (APAF) for the proteomic services, Patrick AM Jansen (Radboud University Medical Center) for the provision of the rabbit anti-*Pf*PanK1 final bleed, Alexander Maier (ANU) for the provision of the pGlux-1 plasmid and the HSP70 antibody, Parichat Prommana (BIOTEC) for the provision of the eGFP-*glmS* plasmid, Régis P Lemaitre (MPI-CBG) for the provision of the DefBac bacmid and Erick Tjhin (ANU) for invaluable advice. XL was supported by an *Australian Government Research Training Program* (AGRTP) scholarship.

## SUPPORTING INFORMATION

**Table S1.**
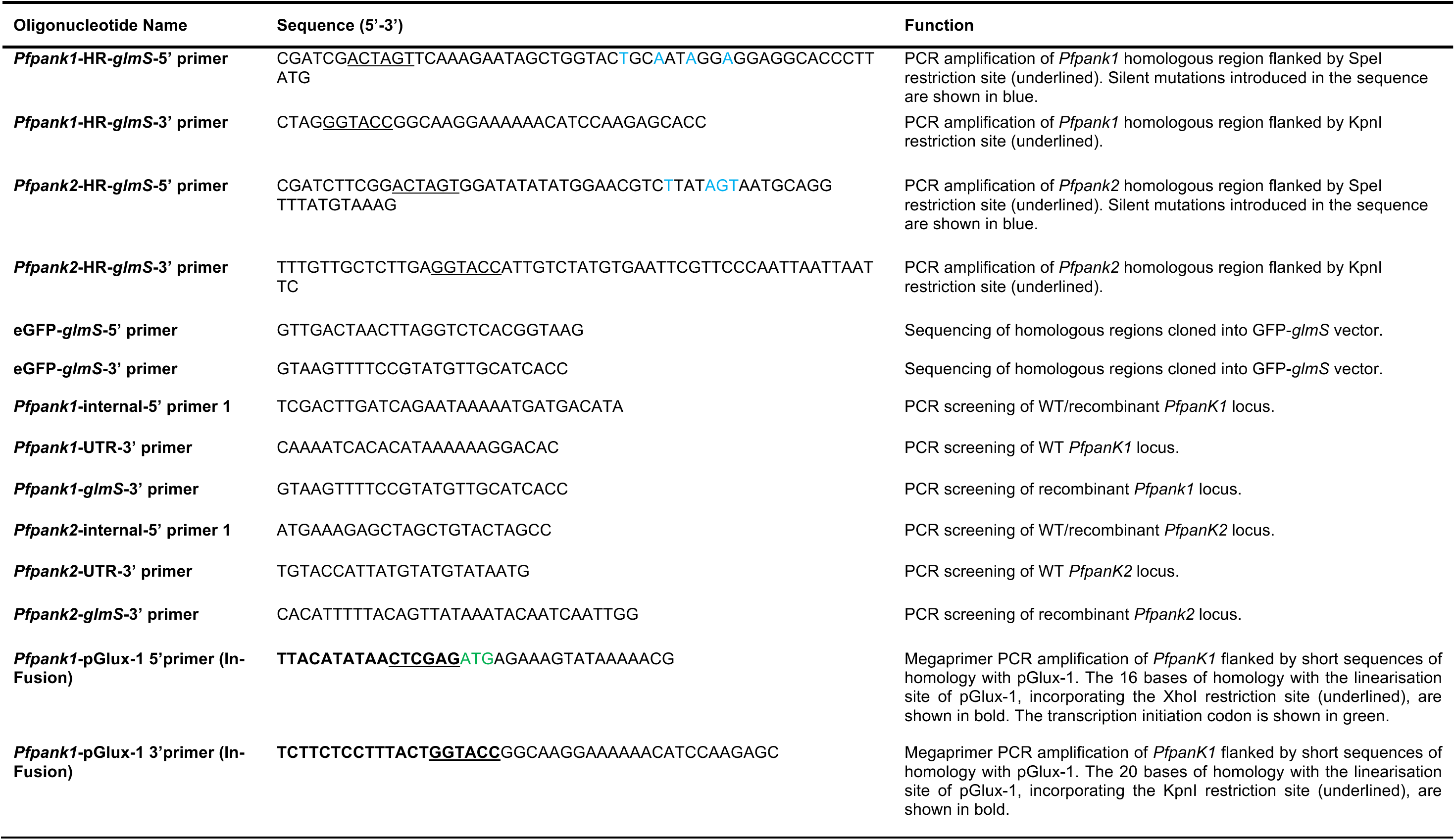
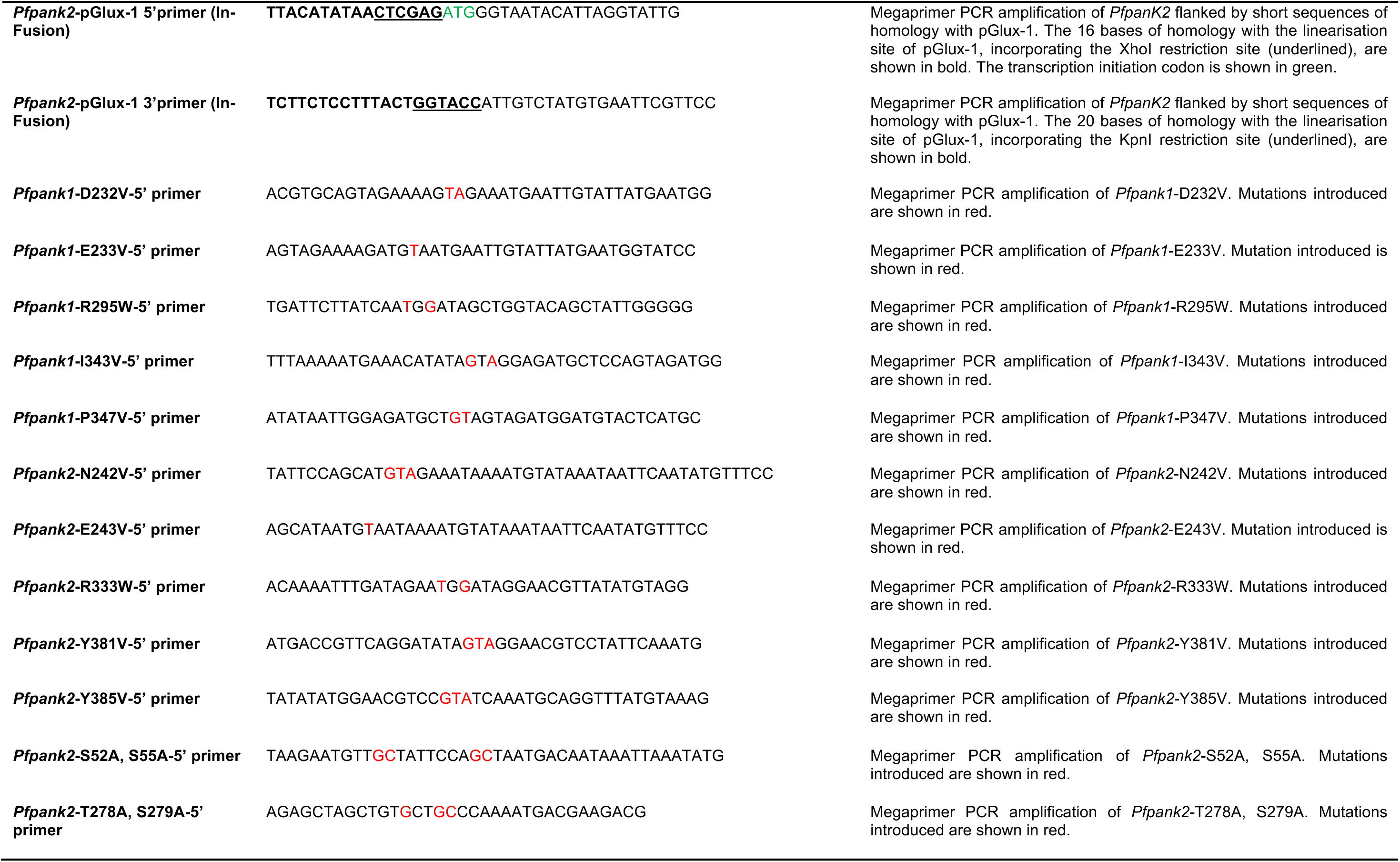
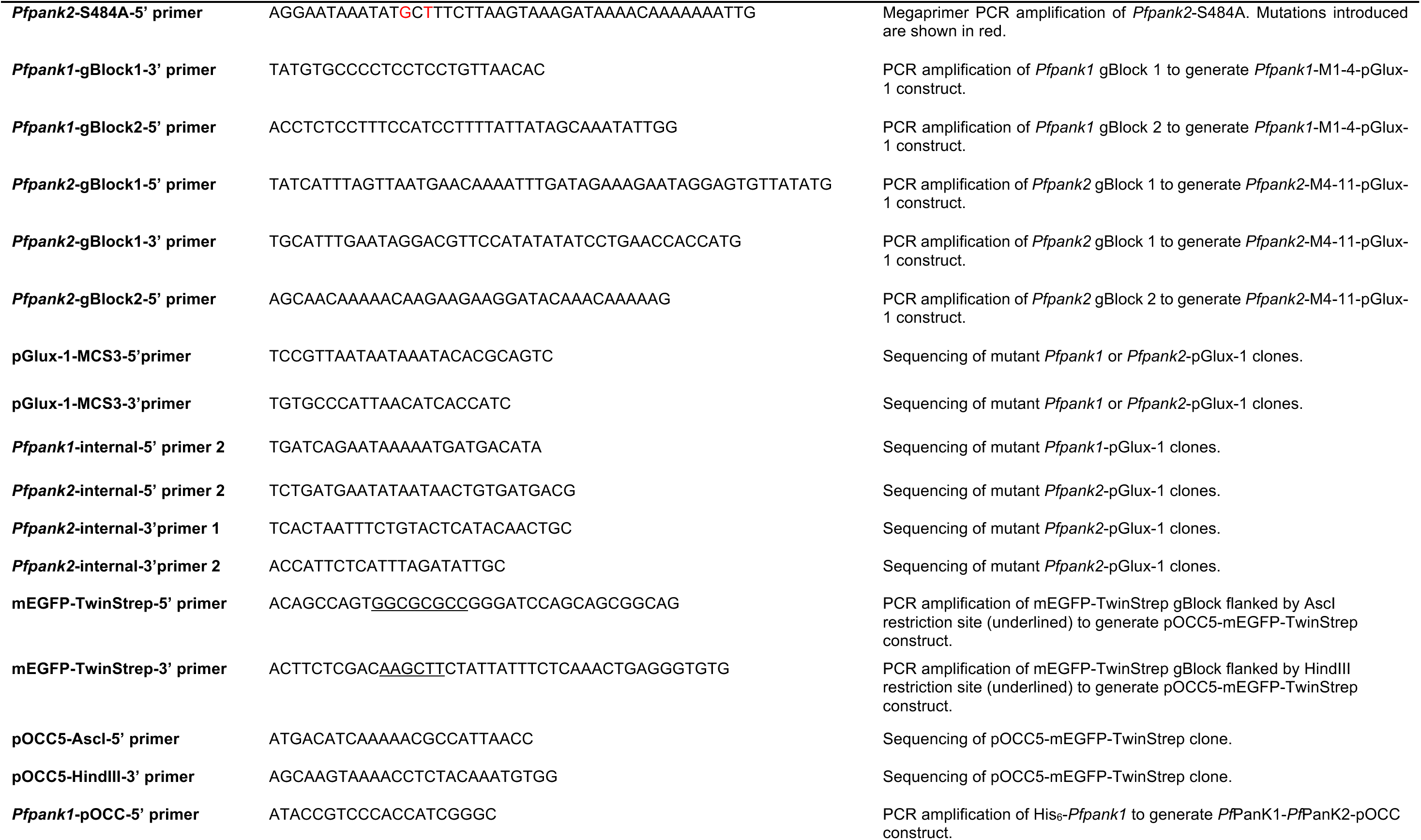
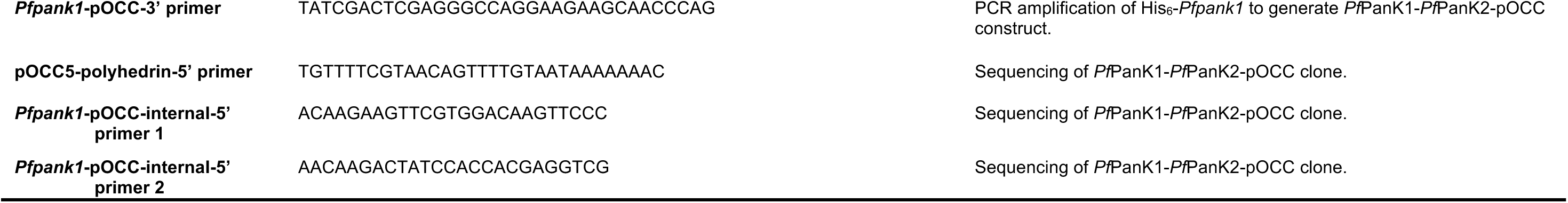
List of oligonucleotides used in this study.

**Table S2.**
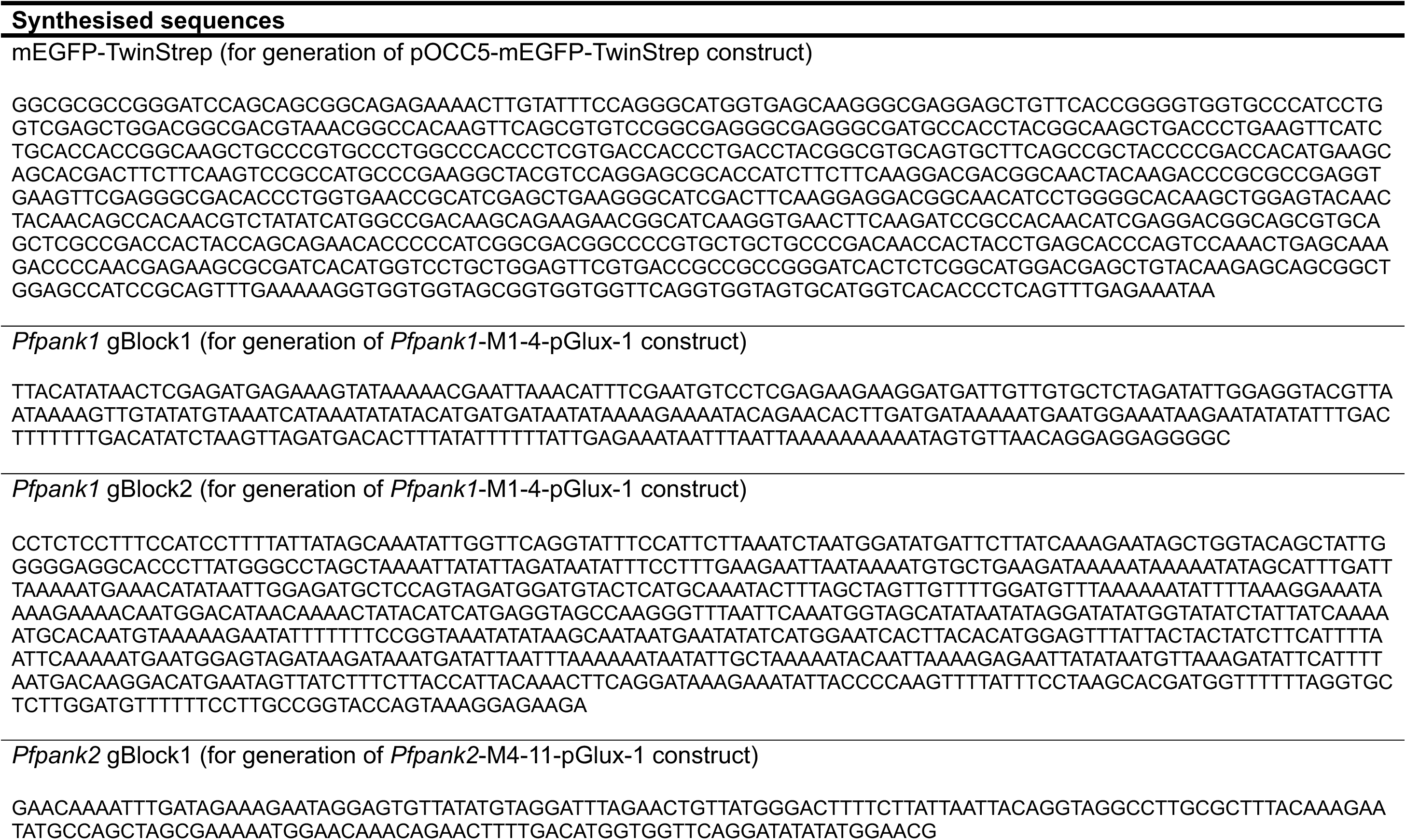
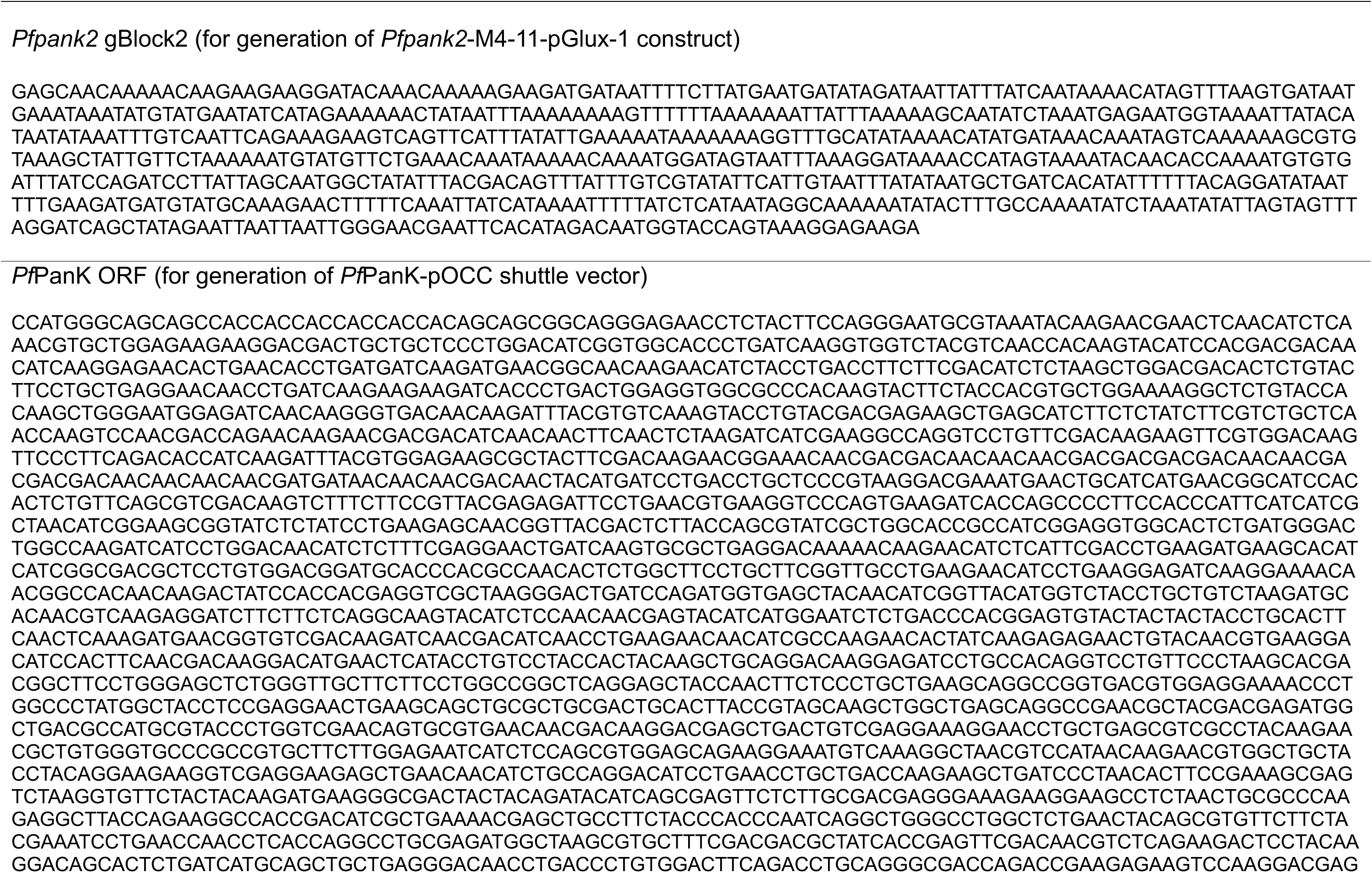
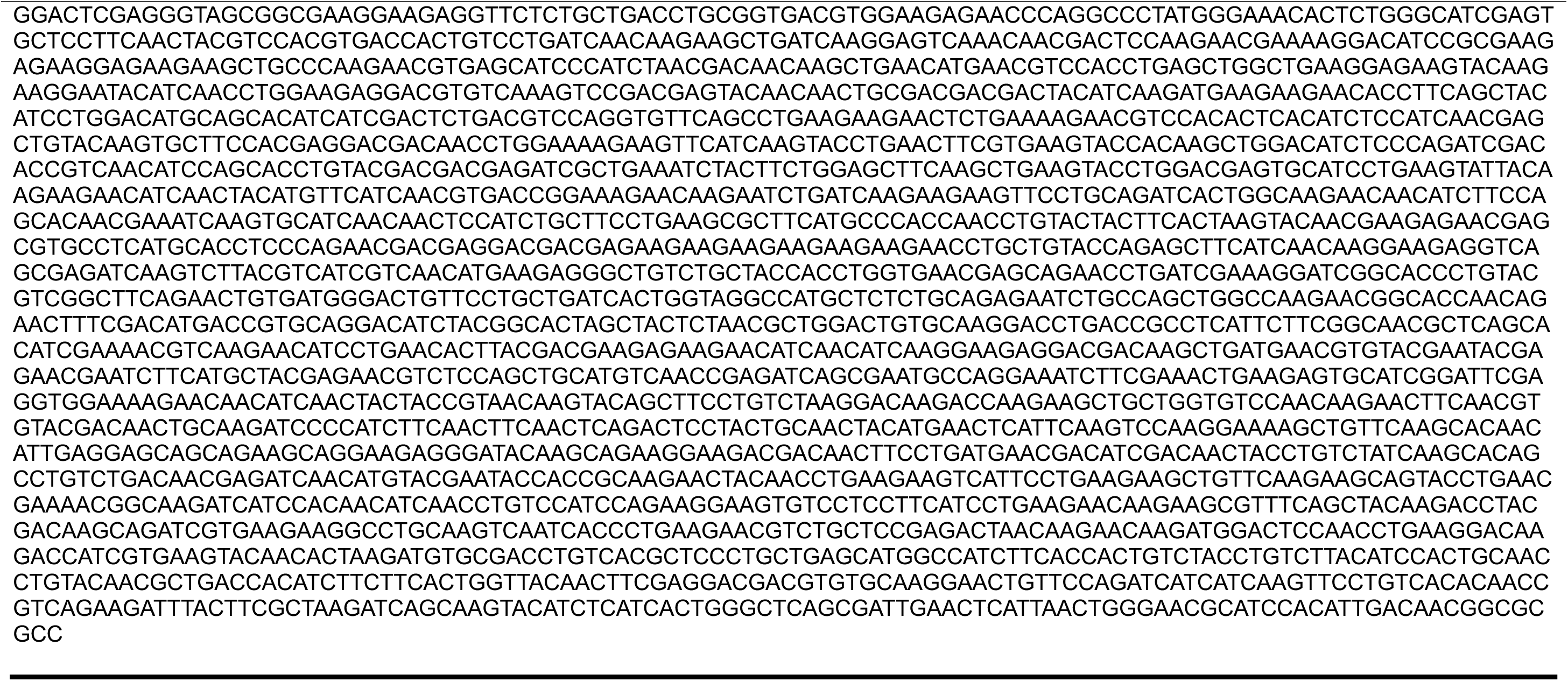
List of synthesised sequences used in this study.

**Figure S1.**
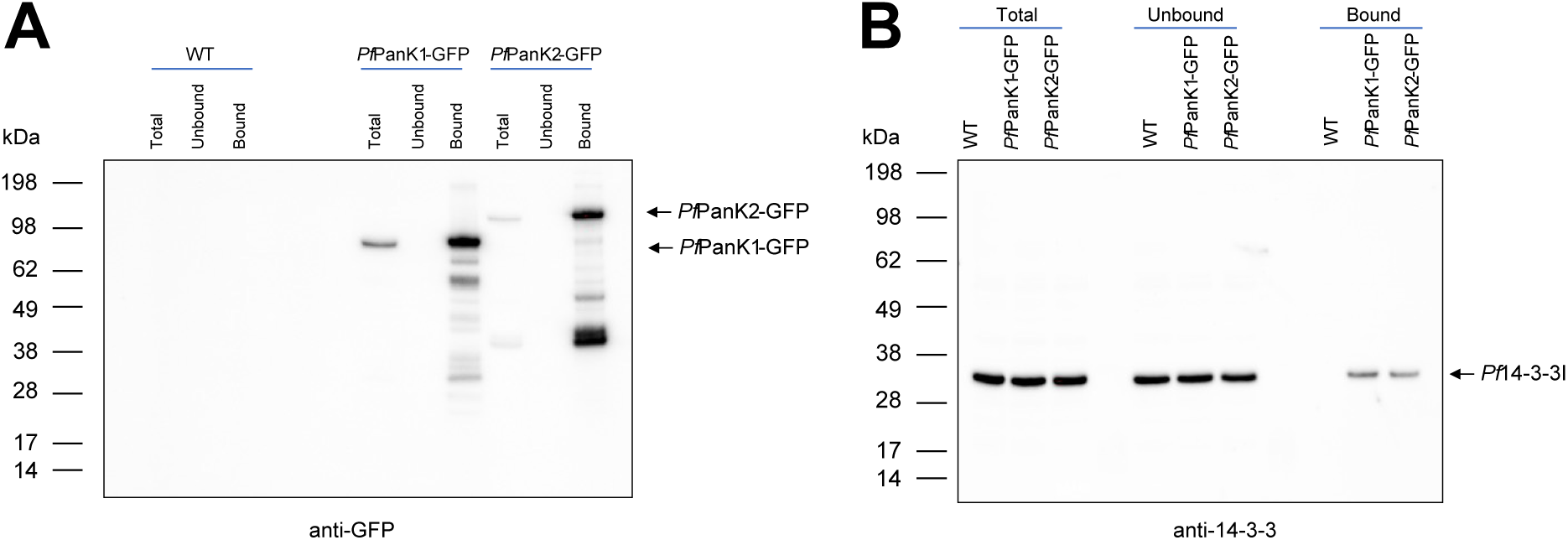
GFP-Trap immunoprecipitation of proteins from WT and transgenic parasites expressing a GFP-tagged *Pf*PanK1 or *Pf*PanK2 under control of the endogenous promoter. Denaturing western blot analysis of the proteins present in the total lysate, unbound and GFP-Trap-bound fractions of *Pf*PanK1-GFP, *Pf*PanK2-GFP and WT lines using anti-GFP **(A)** or pan anti-14-3-3 antibody **(B)**. The western blot shown is a representative of two independent experiments, each performed with a different batch of parasites.

**Figure S2.**
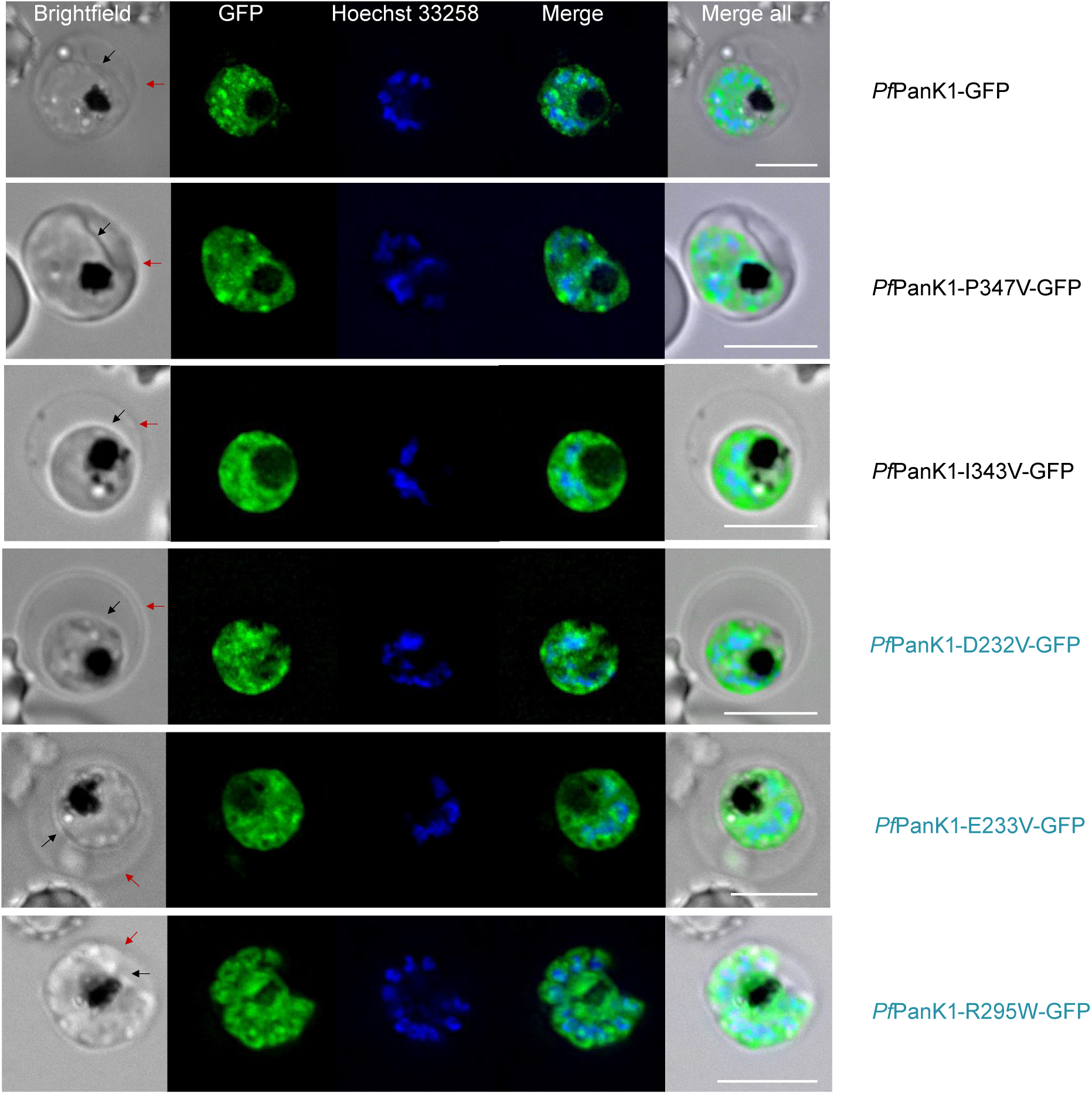
Localisation of mutant *Pf*PanK1-GFP in erythrocytes infected with mutant *Pf*PanK1-GFP transgenic parasites. Nuclei of parasites were stained with Hoechst 33258 (blue). From left to right: brightfield; GFP fluorescence; Hoechst 33258; merge of GFP fluorescence and Hoechst 33258; merge of all three channels. Mutant *Pf*PanK1-GFP fluorescence (green) was observed throughout the cytosol of the transgenic parasites. The intensity of GFP fluorescence was often greatest in regions overlapping with parasite nuclei. The parasite membrane is indicated with a black arrow and the erythrocyte membrane is indicated with a red arrow. Scale bar: 5 μm.

**Figure S3.**
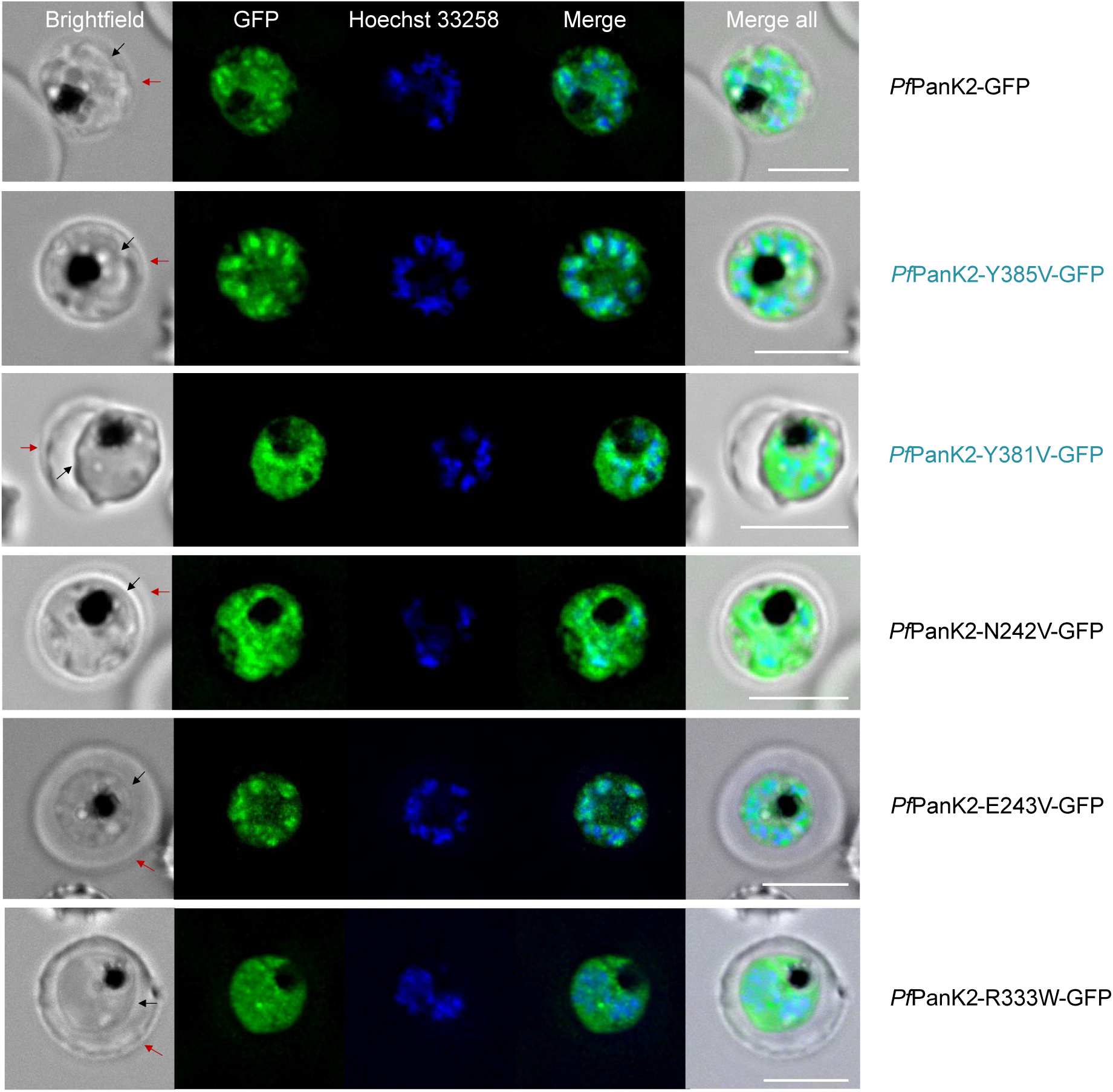
Localisation of mutant *Pf*PanK2-GFP in erythrocytes infected with mutant *Pf*PanK2-GFP transgenic parasites. Nuclei of parasites were stained with Hoechst 33258 (blue). From left to right: brightfield; GFP fluorescence; Hoechst 33258; merge of GFP fluorescence and Hoechst 33258; merge of all three channels. Mutant *Pf*PanK2-GFP fluorescence (green) was observed throughout the cytosol of the transgenic parasites. The intensity of GFP fluorescence was often greatest in regions overlapping with parasite nuclei. The parasite membrane is indicated with a black arrow and the erythrocyte membrane is indicated with a red arrow. Scale bar: 5 μm.

**Figure S4.**
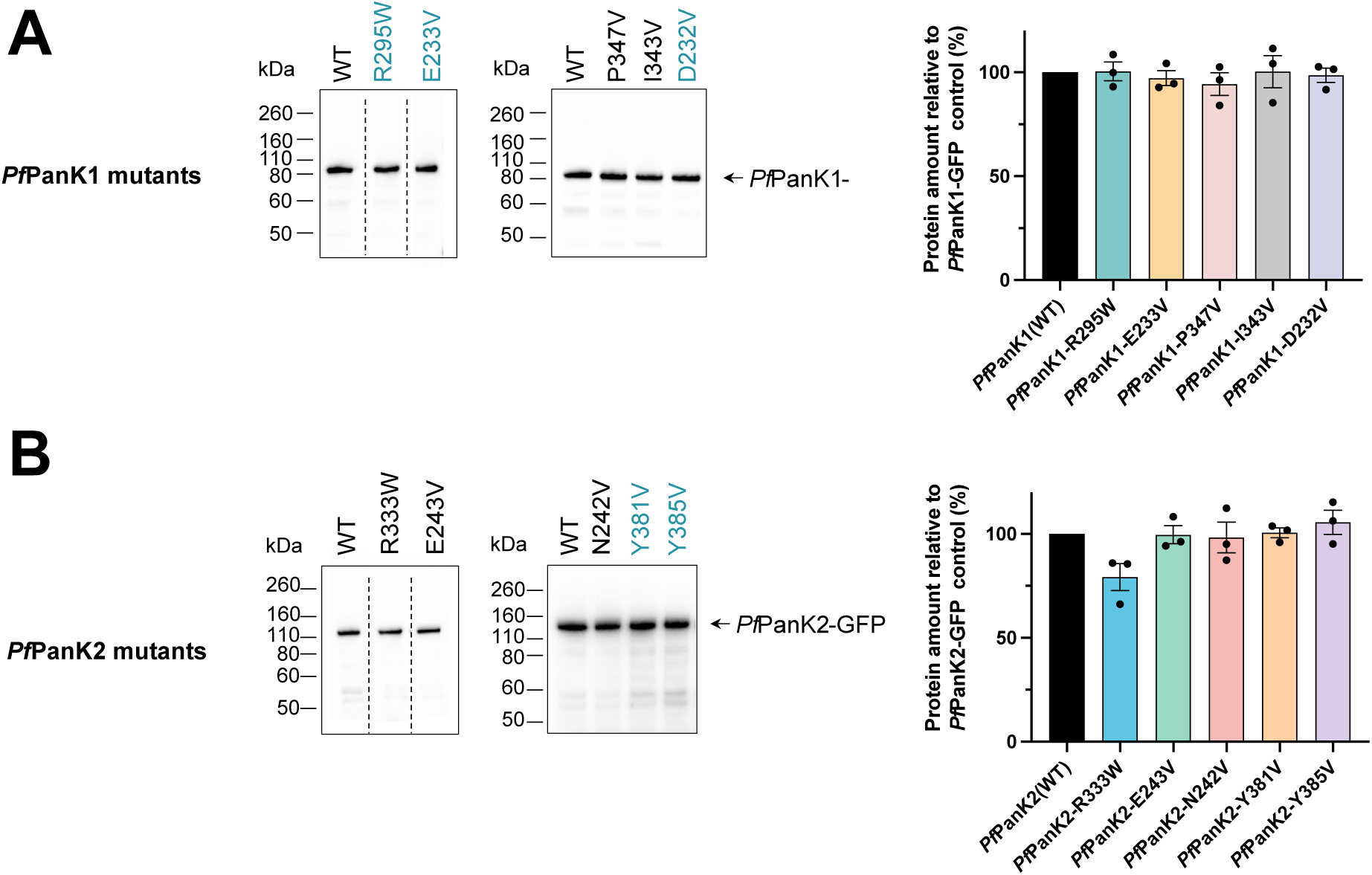
Analysis of GFP-Trap immunoprecipitation of proteins from WT and mutant *Pf*PanK1-GFP (A) or *Pf*PanK2-GFP (B) parasites. Anti-GFP denaturing western blot analysis of the GFP-Trap immunoprecipitated complexes that were used in the [^14^C]pantothenate phosphorylation assays in Figure 3B and **3C**. The western blots shown are representative of three independent experiments, each performed with a different batch of parasites. Dashed lines on the left-hand-side blots indicates where the membranes were cut to remove extraneous sections, but the included lanes within each blot are from the same membrane. Densitometry analysis was performed using Fiji version 2.1.0. Values are presented as a percentage relative to the amount of protein present in the WT samples. Values are averaged from three independent experiments and error bars represent SEM.

**Figure S5.**
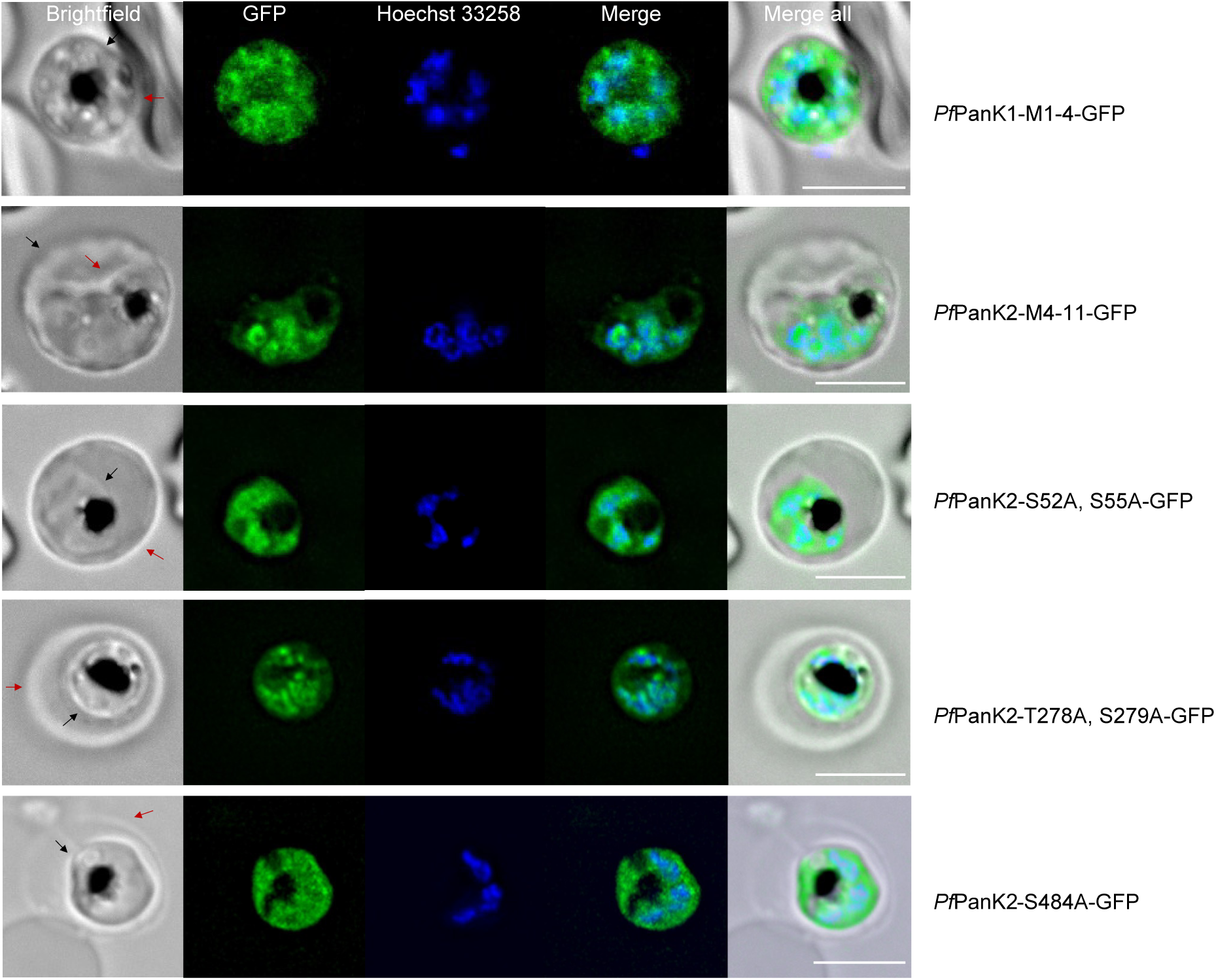
Localisation of *Pf*14-3-3I binding site mutant *Pf*PanK1-GFP or *Pf*PanK2-GFP in erythrocytes infected with mutant *Pf*PanK1-GFP or *Pf*PanK2-GFP transgenic parasites. Nuclei of parasites were stained with Hoechst 33258 (blue). From left to right: brightfield; GFP fluorescence; Hoechst 33258; merge of GFP fluorescence and Hoechst 33258; merge of all three channels. Mutant *Pf*PanK1-GFP or *Pf*PanK2-GFP fluorescence (green) was observed throughout the cytosol of the transgenic parasites. The intensity of GFP fluorescence was often greatest in regions overlapping with parasite nuclei. Parasite membrane is indicated with a black arrow and erythrocyte membrane is indicated with a red arrow. Scale bar: 5 μm.

**Figure S6.**
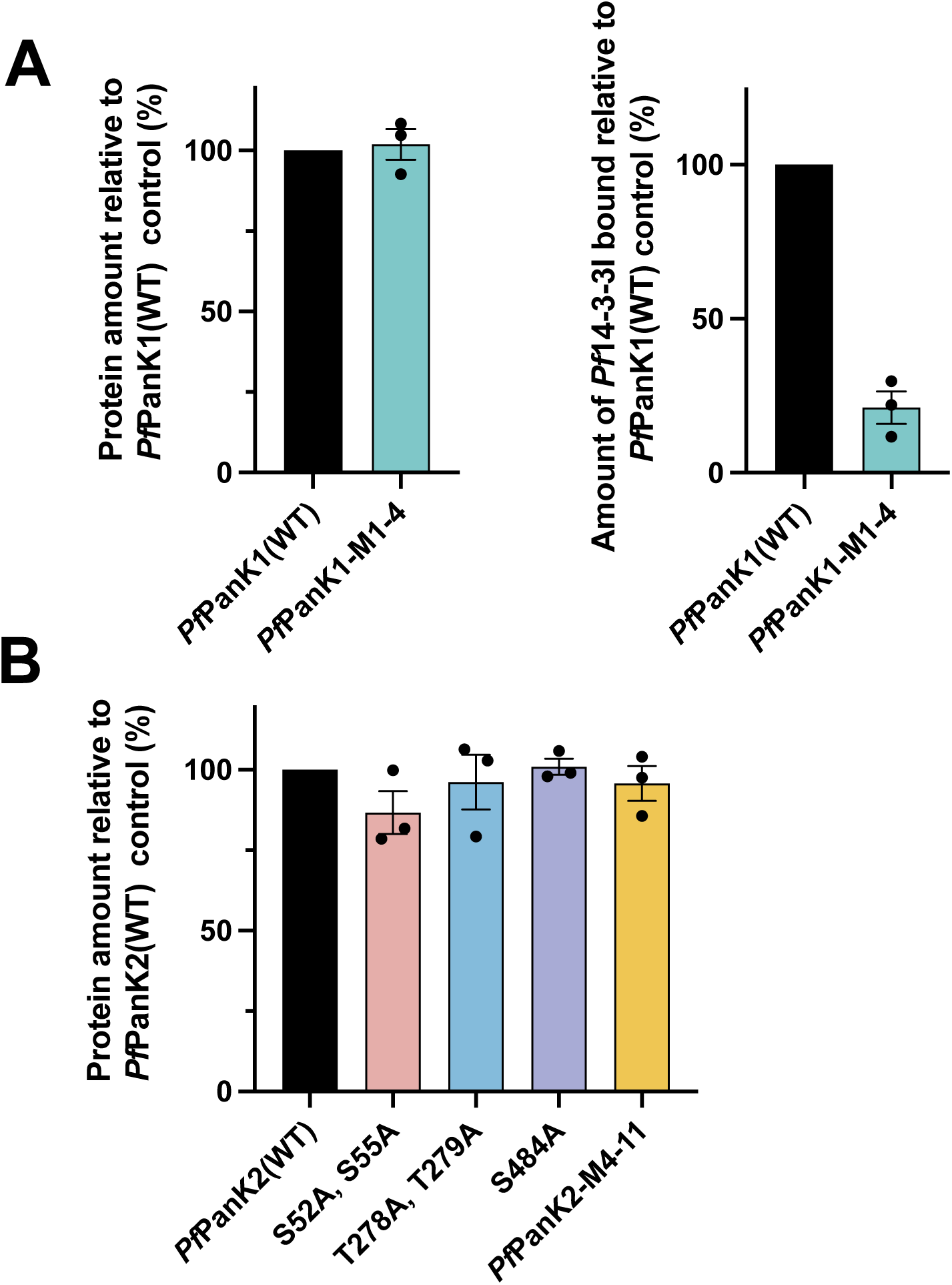
Densitometry analysis of GFP-Trap immunoprecipitation of proteins from WT and putative *Pf*14-3-3I binding site mutant *Pf*PanK1-GFP (A) or *Pf*PanK2-GFP (B) parasites. Densitometry analysis of *Pf*PanK1-GFP (A, left) or *Pf*PanK2-GFP in the GFP-Trap immunoprecipitated complexes that were used in the denaturing western blots and the [^14^C]pantothenate phosphorylation assays in Figure 7B and **7C**. Panel A (right) shows the amount of *Pf*14-3-3I bound to the GFP-Trap immunoprecipitated *Pf*PanK1-M1-4 complex compared to that bound to the *Pf*PanK1-WT complex. Densitometry analysis was performed using Fiji version 2.1.0. Values are presented as a percentage relative to the amount of protein present in the WT samples. Values are averaged from three independent experiments and error bars represent SEM.

**Figure S7.**
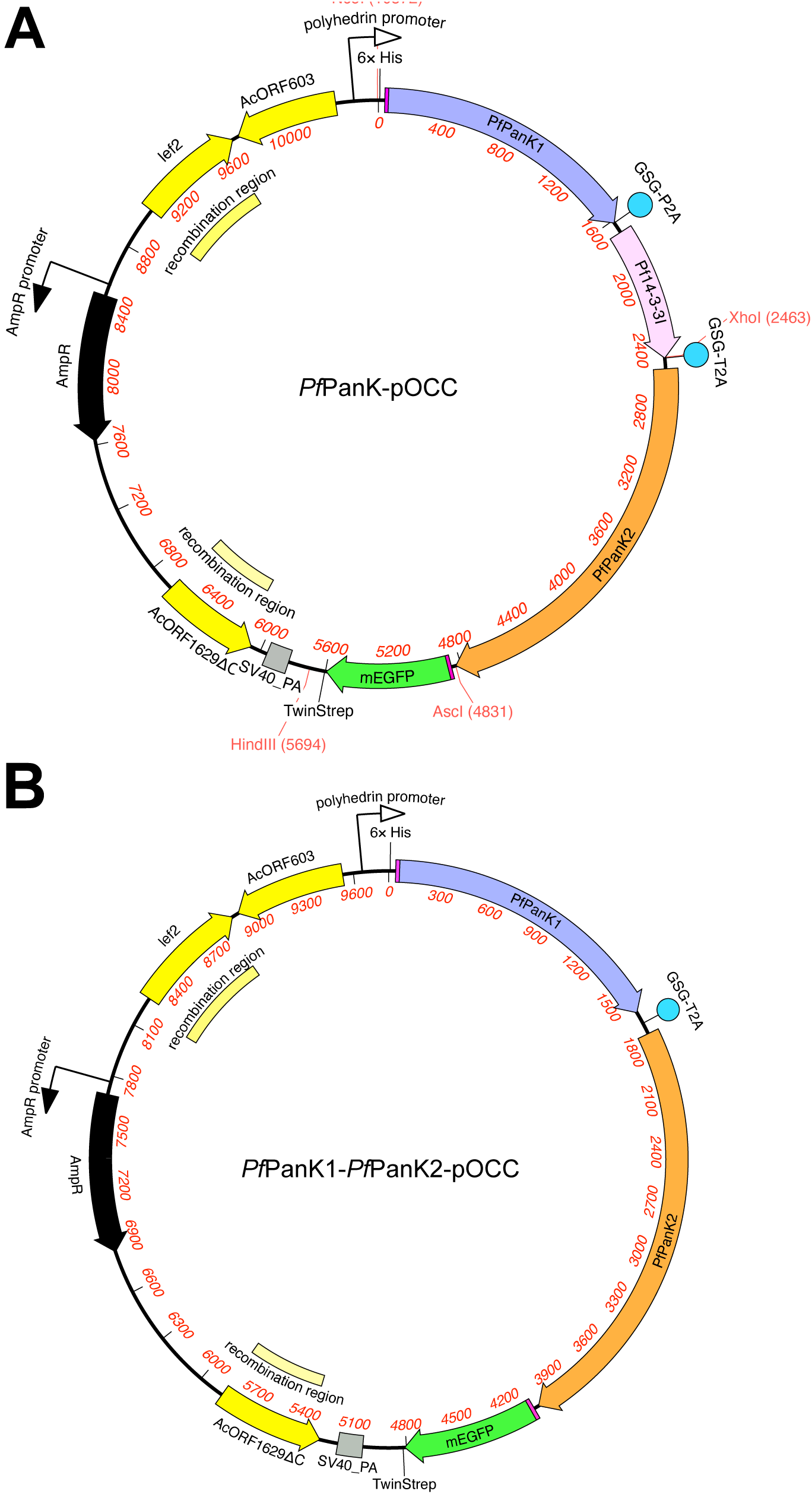
Vector maps of the *Pf*PanK-pOCC (A) and the *Pf*PanK1-*Pf*PanK2-pOCC (B) constructs. The constructs carry the *Pf*PanK or the *Pf*PanK1-PfPanK2 expression cassette downstream of the polyhedron promoter. Two homology regions are present to rescue the doubly defective DefBac bacmid upon infection, ensuring that all infective viruses contain the recombinant target gene. The constructs also bear a 𝛽*-lactamase* (ampicillin resistant, AmpR) gene for positive selection in *E. coli*. SV40_PA: Simian Virus 40 polyadenylation signal. Lef2: complementary *lef2* gene truncation. AcORF1629ΔC: complementary *AcORF1629* gene truncation, AcORF603: full length *AcORF603* gene.

**Figure S8.**
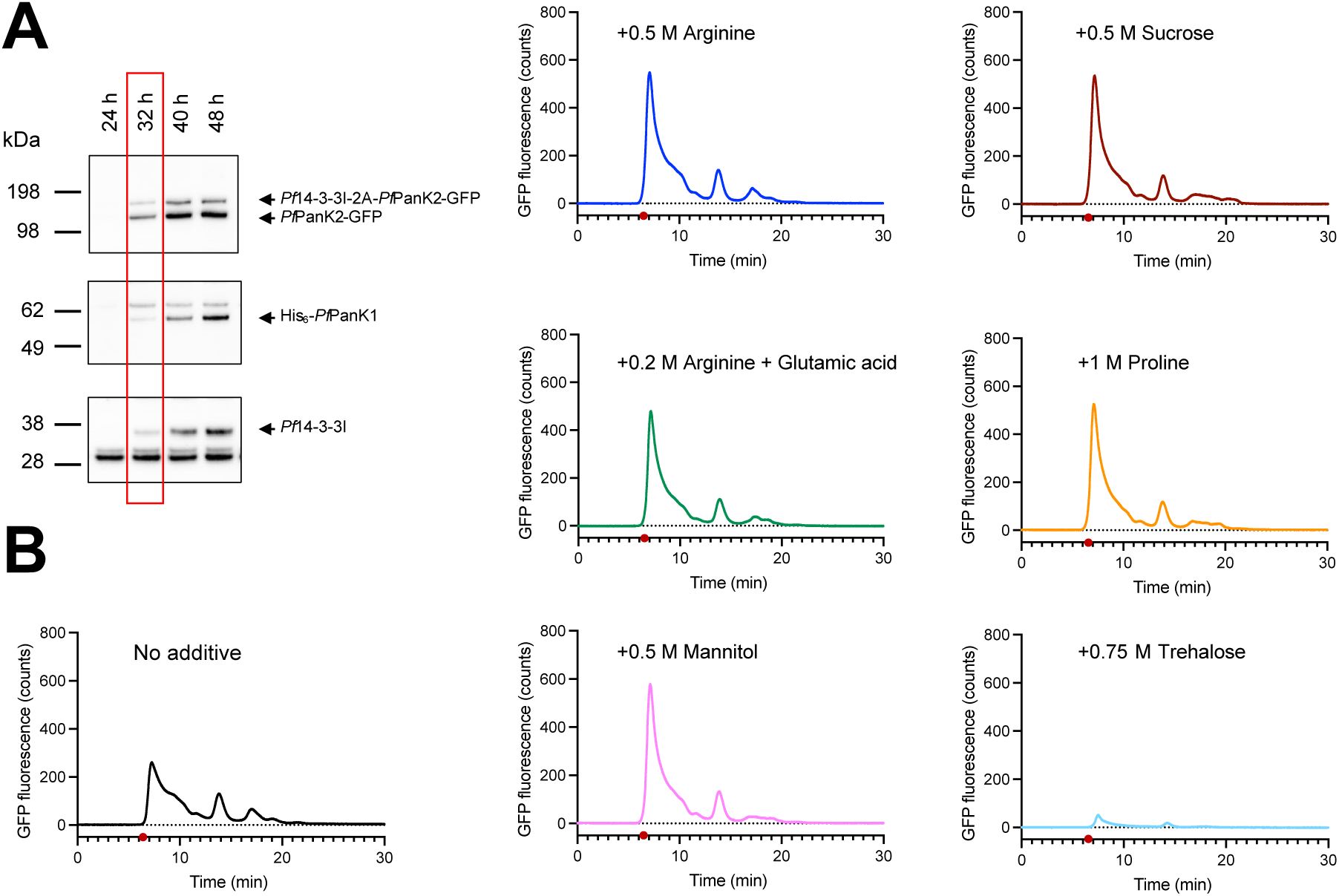
Attempts to optimise the *Pf*PanK complex expression in insect cells. **(A)** A viral infection time course study was performed to optimise expression levels. Samples at each time point were analysed by denaturing western blot to check expression level using anti-GFP, anti-*Pf*PanK1 or pan anti-14-3-3 antibodies. **(B)** FSEC traces of the *Pf*PanK-GFP (with *Pf*14-3-3I) complex expressed in insect cells 32 h post infection. Cells were lysed in 50 mM Tris-HCl, 500 mM NaCl, 10 % glycerol, pH 7.5 with and without the addition of the indicated stabilising additives [59]. None of the additives tested resolved the aggregation issue. Void is marked with a red dot.

**Figure S9.**
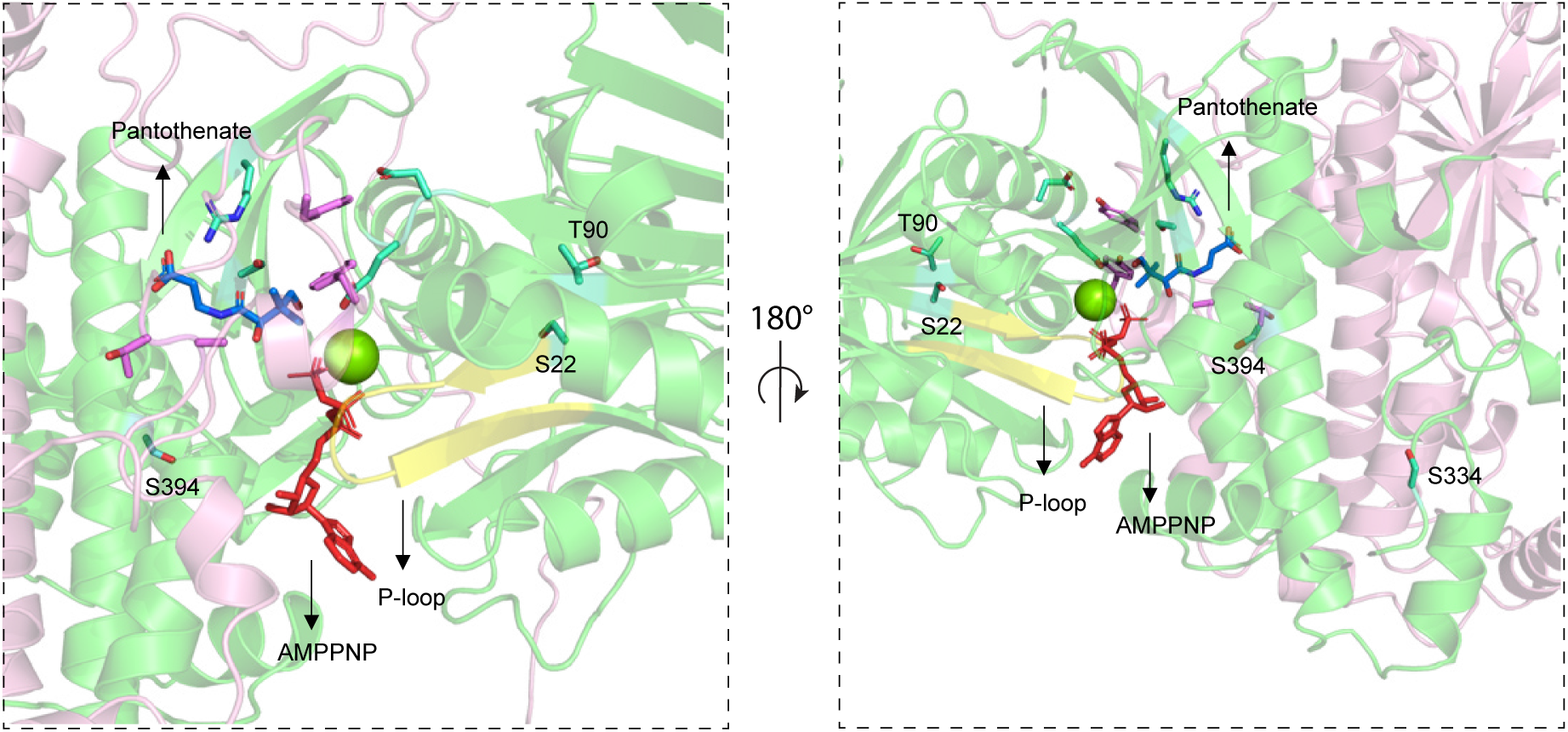
Predicted location of *Pf*PanK1 S22, T90, S334 and S394 in the *Pf*PanK1-*Pf*PanK2 heteromer. The right panel shows the same structure as the left panel, rotated 180° around the Y-axis and shown at reduced magnification. The heteromeric *Pf*PanK is composed of *Pf*PanK1 (green) and *Pf*PanK2 (pink). Side chains of residues from *Pf*PanK1 are shown in cyan and residues from *Pf*PanK2 are shown in pink. The P-loop is shown in yellow, AMPPNP in red, pantothenate in blue and Mg^2+^ as a green ball.

## References

1. World Health Organization (2024) World malaria report 2024 Geneva

2. Dondorp, A. M., Nosten, F., Yi, P., Das, D., Phyo, A. P., Tarning, J. et al. (2009) Artemisinin resistance in *Plasmodium falciparum* malaria N Engl J Med 361, 455–467

3. Fairhurst, R. M., and Dondorp, A. M. (2016) Artemisinin-resistant Plasmodium falciparum malaria Microbiol Spectr 4, EI10–0013-2016

4. Bergmann, C., van Loon, W., Habarugira, F., Tacoli, C., Jager, J. C., Savelsberg, D., et al. (2021) Increase in Kelch 13 Polymorphisms in *Plasmodium falciparum*, Southern Rwanda Emerg Infect Dis 27, 294–296

5. Yoshida, N., Yamauchi, M., Morikawa, R., Hombhanje, F., and Mita, T. (2021) Increase in the proportion of *Plasmodium falciparum* with kelch13 C580Y mutation and decline in pfcrt and pfmdr1 mutant alleles in Papua New Guinea Malar J 20, 410

6. Wells, T. N., Hooz van Huijsduijnen, R., and Van Voorhis, W. C. (2015) Malaria medicines: a glass half full? Nat Rev Drug Discov 14, 424–442

7. Burrows, J. N., Duparc, S., Gutteridge, W. E.,Hooz van Huijsduijnen, R., Kaszubska, W., Macintyre, F. et al. (2017) New developments in anti-malarial target candidate and product profiles Malar J 16, 26

8. Daugherty, M., Polanuyer, B., Farrell, M., Scholle, M., Lykidis, A., de Crecy-Lagard, V., et al. (2002) Complete reconstitution of the human coenzyme A biosynthetic pathway via comparative genomics J Biol Chem 277, 21431–21439

9. Leonardi, R., Zhang, Y. M., Rock, C. O., and Jackowski, S. (2005) Coenzyme A: back in action Prog Lipid Res 44, 125–153

10. Strauss, E. (2010) Coenzyme A Biosynthesis and Enzymology In Comprehensive natural products II - Chemistry and biology, Liu; H-W, and Mander L, eds. Elsevier, 315–410

11. de Vries, L. E., Jansen, P. A. M., Barcelo, C., Munro, J., Verhoef, J. M. J., Pasaje, C. F. A. et al. (2022) Preclinical characterization and target validation of the antimalarial pantothenamide MMV693183 Nat Commun 13, 2158

12. de Vries, L. E., Lunghi, M., Krishnan, A., Kooij, T. W. A., and Soldati-Favre, D. (2021) Pantothenate and CoA biosynthesis in Apicomplexa and their promise as antiparasitic drug targets PLoS Pathog 17, e1010124

13. Guan, J., Spry, C., Tjhin, E. T., Yang, P., Kiàkool, T., Howieson, V. M. et al. (2021) Exploring heteroaromatic rings as a replacement for the labile amide of antiplasmodial pantothenamides J Med Chem 64, 4478–4497

14. Liu, X., Thistlethwaite, S., Kholiya, R., Pierscianowski, J., Saliba, K. J., and Auclair, K. (2024) Chemical synthesis and enzymatic late-stage diversification of novel pantothenate analogues with antiplasmodial activity Eur J Med Chem 280, 116902

15. Schalkwijk, J., Allman, E. L., Jansen, P. A. M., de Vries, L. E., Verhoef, J. M. J., Jackowski, S., et al. (2019) Antimalarial pantothenamide metabolites target acetyl-coenzyme A biosynthesis in Plasmodium falciparum Sci Transl Med 11, eaas9917

16. Spry, C., Kirk, K., and Saliba, K. J. (2008) Coenzyme A biosynthesis: an antimicrobial drug target FEMS Microbiol Rev 32, 56–106

17. Jackowski, S., and Rock, C. O. (1981) Regulation of coenzyme A biosynthesis J Bacteriol 148, 926–932

18. Rock, C. O., Calder, R. B., Karim, M. A., and Jackowski, S. (2000) Pantothenate kinase regulation of the intracellular concentration of coenzyme A J Biol Chem 275, 1377–1383

19. Lehane, A. M., Marcheà, R. V., Spry, C., van Schalkwyk, D. A., Teng, R., Kirk, K. et al. (2007) Feedback inhibition of pantothenate kinase regulates pantothenol uptake by the malaria parasite J Biol Chem 282, 25395–25405

20. Tjhin, E. T., Spry, C., Sewell, A. L., Hoegl, A., Barnard, L., Sexton, A. E., et al. (2018) Mutations in the pantothenate kinase of *Plasmodium falciparum* confer diverse sensitivity profiles to antiplasmodial pantothenate analogues PLoS Pathog 14, e1006918

21. Brand, L. A., and Strauss, E. (2005) Characterization of a new pantothenate kinase isoform from *Helicobacter pylori* J Biol Chem 280, 20185–20188

22. Song, W. J., and Jackowski, S. (1992) Cloning, sequencing, and expression of the pantothenate kinase (coaA) gene of *Escherichia coli* J Bacteriol 174, 6411–6417

23. Calder, R. B., Williams, R. S., Ramaswamy, G., Rock, C. O., Campbell, E., Unkles, S. E. et al. (1999) Cloning and characterization of a eukaryotic pantothenate kinase gene (panK) from *Aspergillus nidulans* J Biol Chem 274, 2014–2020

24. Yun, M., Park, C. G., Kim, J. Y., Rock, C. O., Jackowski, S., and Park, H. W. (2000) Structural basis for the feedback regulation of *Escherichia coli* pantothenate kinase by coenzyme A J Biol Chem 275, 28093–28099

25. Das, S., Kumar, P., Bhor, V., Surolia, A., and Vijayan, M. (2006) Invariance and variability in bacterial PanK: a study based on the crystal structure of *Mycobacterium tuberculosis* PanK *Acta Crystallogr* D Biol Crystallogr 62, 628–638

26. Yang, K., Eyobo, Y., Brand, L. A., Martynowski, D., Tomchick, D., Strauss, E. et al. (2006) Crystal structure of a type III pantothenate kinase: insight into the mechanism of an essential coenzyme A biosynthetic enzyme universally distributed in bacteria J Bacteriol 188, 5532–5540

27. Hong, B. S., Yun, M. K., Zhang, Y. M., Chohnan, S., Rock, C. O., White, S. W. et al. (2006) Prokaryotic type II and type III pantothenate kinases: The same monomer fold creates dimers with distinct catalytic properties Structure 14, 1251–1261

28. Nicely, N. I., Parsonage, D., Paige, C., Newton, G. L., Fahey, R. C., Leonardi, R. et al. (2007) Structure of the type III pantothenate kinase from *Bacillus anthracis* at 2.0 A resolution: implications for coenzyme A-dependent redox biology Biochemistry 46, 3234–3245

29. Hong, B. S., Senisterra, G., Rabeh, W. M., Vedadi, M., Leonardi, R., Zhang, Y. M. et al. (2007) Crystal structures of human pantothenate kinases. Insights into allosteric regulation and mutations linked to a neurodegeneration disorder J Biol Chem 282, 27984–27993

30. Li, B., Tempel, W., Smil, D., Bolshan, Y., Schapira, M., and Park, H. W. (2013) Crystal structures of *Klebsiella pneumoniae* pantothenate kinase in complex with N-substituted pantothenamides Proteins 81, 1466–1472

31. Franklin, M. C., Cheung, J., Rudolph, M. J., Burshteyn, F., Cassidy, M., Gary, E. et al. (2015) Structural genomics for drug design against the pathogen Proteins-Structure Function and Bioinformatics 83, 2124–2136

32. Spry, C., van Schalkwyk, D. A., Strauss, E., and Saliba, K. J. (2010) Pantothenate utilization by *Plasmodium* as a target for antimalarial chemotherapy Infect Disord Drug Targets 10, 200–216

33. Tjhin, E. T., Howieson, V. M., Spry, C., van Dooren, G. G., and Saliba, K. J. (2021) A novel heteromeric pantothenate kinase complex in apicomplexan parasites PLoS Pathog 17, e1009797

34. Zhang, M., Wang, C., Otto, T. D., Oberstaller, J., Liao, X., Adapa, S. R. et al. (2018) Uncovering the essential genes of the human malaria parasite *Plasmodium falciparum* by saturation mutagenesis Science 360, eaap7847

35. Lalle, M., Curra, C., Ciccarone, F., Pace, T., Ceccheà, S., Fantozzi, L. et al. (2011) Dematin, a component of the erythrocyte membrane skeleton, is internalized by the malaria parasite and associates with *Plasmodium* 14-3-3 J Biol Chem 286, 1227–1236

36. Sluchanko, N. N., and Gusev, N. B. (2017) Moonlighting chaperone-like activity of the universal regulatory 14-3-3 proteins FEBS J 284, 1279–1295

37. van Hemert, M. J., Steensma, H. Y., and van Heusden, G. P. (2001) 14-3-3 proteins: key regulators of cell division, signalling and apoptosis Bioessays 23, 936–946

38. Muslin, A. J., and Xing, H. (2000) 14-3-3 proteins: regulation of subcellular localization by molecular interference Cell Signal 12, 703–709

39. Bridges, D., and Moorhead, G. B. (2005) 14-3-3 proteins: a number of functions for a numbered protein Sci STKE 2005, re10

40. Yaffe, M. B. (2002) How do 14-3-3 proteins work?-- Gatekeeper phosphorylation and the molecular anvil hypothesis FEBS LeN 513, 53–57

41. Sluchanko, N. N., Artemova, N. V., Sudnitsyna, M. V., Safenkova, I. V., Antson, A. A., Levitsky, D. I. et al. (2012) Monomeric 14-3-3zeta has a chaperone-like activity and is stabilized by phosphorylated HspB6 Biochemistry 51, 6127–6138

42. Subramanian, C., Yun, M. K., Yao, J., Sharma, L. K., Lee, R. E., White, S. W. et al. (2016) Allosteric regulation of mammalian pantothenate kinase J Biol Chem 291, 22302–22314

43. Divo, A. A., Geary, T. G., Davis, N. L., and Jensen, J. B. (1985) Nutritional retiuirements of *Plasmodium falciparum* in culture. I. Exogenously supplied dialyzable components necessary for continuous growth J Protozool 32, 59–64

44. Yaffe, M. B., Riànger, K., Volinia, S., Caron, P. R., Aitken, A., Leffers, H. et al. (1997) The structural basis for 14-3-3:phosphopeptide binding specificity Cell 91, 961–971

45. Treeck, M., Sanders, J. L., Elias, J. E., and Boothroyd, J. C. (2011) The phosphoproteomes of *Plasmodium falciparum* and *Toxoplasma gondii* reveal unusual adaptations within and beyond the parasites’ boundaries Cell Host Microbe 10, 410–419

46. Solyakov, L., Halbert, J., Alam, M. M., Semblat, J. P., Dorin-Semblat, D., Reininger, L. et al. (2011) Global kinomic and phospho-proteomic analyses of the human malaria parasite *Plasmodium falciparum* Nat Commun 2, 565

47. Lasonder, E., Green, J. L., Camarda, G., Talabani, H., Holder, A. A., Langsley, G. et al. (2012) The Schizont Phosphoproteome Reveals Extensive Phosphatidylinositol and cAMP-Protein Kinase A Signaling J Proteome Res 11, 5323–5337

48. Pease, B. N., Huttlin, E. L., Jedrychowski, M. P., Talevich, E., Harmon, J., Dillman, T. et al. (2013) Global analysis of protein expression and phosphorylation of three stages of *Plasmodium falciparum* intraerythrocytic development J Proteome Res 12, 4028–4045

49. Lasonder, E., Green, J. L., Grainger, M., Langsley, G., and Holder, A. A. (2015) Extensive differential protein phosphorylation as intraerythrocytic schizonts develop into extracellular invasive merozoites Proteomics 15, 2716–2729

50. Tjhin, E. T. (2018) The pantothenate kinase of the human malaria parasite Plasmodium falciparum, Australian National University

51. Dastidar, E. G., Dzeyk, K., Krijgsveld, J., Malmtiuist, N. A., Doerig, C., Scherf, A. et al. (2013) Comprehensive histone phosphorylation analysis and identification of *Pf*14-3-3 protein as a histone H3 phosphorylation reader in malaria parasites Plos One 8,

52. Jain, R., Dey, P., Gupta, S., Pati, S., Bhaxacherjee, A., Munde, M. et al. (2020) Molecular dynamics simulations and biochemical characterization of 14-3-3 and CDPK1 interaction towards its role in growth of human malaria parasite Biochem J 477, 2153–2177

53. More, K. R., Kaur, I., Giai Gianetto, Q., Invergo, B. M., Chaze, T., Jain, R. et al. (2020) Phosphorylation-dependent assembly of a 14-3-3 mediated signaling complex during red blood cell Invasion by *Plasmodium falciparum* merozoites mBio 11, e01287–01220

54. Sharma, M., Krishnan, D., Singh, A., Negi, P., Rani, K., Revikumar, A. et al. (2025) raf kinase inhibitor is a lipid binding protein that interacts with and regulates the activity of *Pf*CDPK1, an essential plant-like kinase required for red blood cell invasion Biochem Bioph Res Co 749,

55. Jarvis, D. L. (2009) Baculovirus-insect cell expression systems Methods Enzymol 463, 191–222

56. Unger, T., and Peleg, Y. (2012) Recombinant protein expression in the baculovirus-infected insect cell system Methods Mol Biol 800, 187–199

57. Lemaitre, R. P., Bogdanova, A., Borgonovo, B., Woodruff, J. B., and Drechsel, D. N. (2019) FlexiBAC: a versatile, open-source baculovirus vector system for protein expression, secretion, and proteolytic processing BMC Biotechnol 19, 20

58. Minskaia, E., Nicholson, J., and Ryan, M. D. (2013) Optimisation of the foot-and-mouth disease virus 2A co-expression system for biomedical applications BMC Biotechnol 13, 67

59. Ahier, A., and Jarriault, S. (2014) Simultaneous expression of multiple proteins under a single promoter in *CaenorhabdiMs elegans* via a versatile 2A-based toolkit GeneMcs 196, 605-613

60. Brading, R. L., Abbott, W. M., Green, I., Davies, A., and McCall, E. J. (2012) Co-expression of protein phosphatases in insect cells affects phosphorylation status and expression levels of proteins Protein Expr Purif 83, 217–225

61. Abdulrahman, W., Radu, L., Garzoni, F., Kolesnikova, O., Gupta, K., Osz-Papai, J. et al. (2015) The production of multiprotein complexes in insect cells using the baculovirus expression system In Structural Proteomics: High-Throughput Methods, 2 Ed. Owens RJ, ed. Springer New York, New York, NY 91--114

62. Leibly, D. J., Nguyen, T. N., Kao, L. T., Hewitt, S. N., Barrett, L. K., and Van Voorhis, W. C. (2012) Stabilizing additives added during cell lysis aid in the solubilization of recombinant proteins Plos One 7, e52482

63. Lunghi, M., Kloehn, J., Krishnan, A., Varesio, E., Vadas, O., and Soldati-Favre, D. (2022) Pantothenate biosynthesis is critical for chronic infection by the neurotropic parasite Toxoplasma gondii Nat Commun 13, 345

64. Yao, J., Subramanian, C., Rock, C. O., and Jackowski, S. (2019) Human pantothenate kinase 4 is a pseudo-pantothenate kinase Protein Sci 28, 1031–1047

65. Hart, R. J., Cornillot, E., Abraham, A., Molina, E., Nation, C. S., Ben Mamoun, C. et al. (2016) Genetic characterization of *Plasmodium* putative pantothenate kinase genes reveals their essential role in malaria parasite transmission to the mosquito Sci Rep 6, 33518

66. Srivastava, A., Philip, N., Hughes, K. R., Georgiou, K., MacRae, J. I., Barrett, M. P. et al. (2016) Stage-specific changes in *Plasmodium* metabolism required for differentiation and adaptation to different host and vector environments PLoS Pathog 12, e1006094

67. Bushell, E., Gomes, A. R., Sanderson, T., Anar, B., Girling, G., Herd, C. et al. (2017) Functional profiling of a *Plasmodium* genome reveals an abundance of essential genes Cell 170, 260–272 e268

68. Cromer, D., Evans, K. J., Schofield, L., and Davenport, M. P. (2006) Preferential invasion of reticulocytes during late-stage *Plasmodium berghei* infection accounts for reduced circulating reticulocyte levels Int J Parasitol 36, 1389–1397

69. Martin-Jaular, L., Elizalde-Torrent, A., Thomson-Luque, R., Ferrer, M., Segovia, J. C., Herreros-Aviles, E. et al. (2013) Reticulocyte-prone malaria parasites predominantly invade CD71hi immature cells: implications for the development of an in vitro culture for *Plasmodium vivax* Malar J 12, 434

70. Tzivion, G., Luo, Z., and Avruch, J. (1998) A dimeric 14-3-3 protein is an essential cofactor for Raf kinase activity Nature 394, 88–92

71. Zeng, Y., Forbes, K. C., Wu, Z., Moreno, S., Piwnica-Worms, H., and Enoch, T. (1998) Replication checkpoint requires phosphorylation of the phosphatase Cdc25 by Cds1 or Chk1 Nature 395, 507–510

72. Obsil, T., Ghirlando, R., Anderson, D. E., Hickman, A. B., and Dyda, F. (2003) Two 14-3-3 binding motifs are required for stable association of Forkhead transcription factor FOXO4 with 14-3-3 proteins and inhibition of DNA binding Biochemistry 42, 15264–15272

73. Saurin, A. T., Durgan, J., Cameron, A. J., Faisal, A., Marber, M. S., and Parker, P. J. (2008) The regulated assembly of a PKCepsilon complex controls the completion of cytokinesis Nat Cell Biol 10, 891–901

74. Woodcock, J. M., Murphy, J., Stomski, F. C., Berndt, M. C., and Lopez, A. F. (2003) The dimeric versus monomeric status of 14-3-3zeta is controlled by phosphorylation of Ser58 at the dimer interface J Biol Chem 278, 36323–36327

75. Kumagai, A., and Dunphy, W. G. (1999) Binding of 14-3-3 proteins and nuclear export control the intracellular localization of the mitotic inducer Cdc25 Genes Dev 13, 1067–1072

76. Yang, J., Winkler, K., Yoshida, M., and Kornbluth, S. (1999) Maintenance of G2 arrest in the *Xenopus oocyte*: a role for 14-3-3-mediated inhibition of Cdc25 nuclear import EMBO J 18, 2174–2183

77. Shimada, T., Fournier, A. E., and Yamagata, K. (2013) Neuroprotective function of 14-3-3 proteins in neurodegeneration Biomed Res Int 2013, 564534

78. Kundu, A., Shelar, S., Ghosh, A. P., Ballestas, M., Kirkman, R., Nam, H. et al. (2020) 14-3-3 proteins protect AMPK-phosphorylated ten-eleven translocation-2 (TET2) from PP2A-mediated dephosphorylation J Biol Chem 295, 1754–1766

79. Liu, TI., Yang, H., Luo, J., Peng, C., Wang, K., Zhang, G. et al. (2024) 14-3-3 protein augments the protein stability of phosphorylated spastin and promotes the recovery of spinal cord injury through its agonist intervention Elife 12, RP90184

80. Trager, W., and Jensen, J. B. (1976) Human malaria parasites in continuous culture Science 193, 673–675

81. Allen, R. J., and Kirk, K. (2010) *Plasmodium falciparum* culture: the benefits of shaking Mol Biochem Parasitol 169, 63–65

82. Xu, Z., Colosimo, A., and Gruenert, D. C. (2003) Site-directed mutagenesis using the megaprimer method Methods Mol Biol 235, 203–207

83. Leyton, D. L., Sevastsyanovich, Y. R., Browning, D. F., Rossiter, A. E., Wells, T. J., Fitzpatrick, R. E. et al. (2011) Size and conformation limits to secretion of disulfide-bonded loops in autotransporter proteins J Biol Chem 286, 42283–42291

84. Tjhin, E. T., Staines, H. M., van Schalkwyk, D. A., Krishna, S., and Saliba, K. J. (2013) Studies with the *Plasmodium falciparum* hexokinase reveal that *Pf*HT limits the rate of glucose entry into glycolysis FEBS LeN 587, 3182–3187

85. Prommana, P., Uthaipibull, C., Wongsombat, C., Kamchonwongpaisan, S., Yuthavong, Y., Knuepfer, E. et al. (2013) Inducible knockdown of *Plasmodium* gene expression using the glmS ribozyme Plos One 8, e73783

86. Wu, Y., Sifri, C. D., Lei, H. H., Su, X. Z., and Wellems, T. E. (1995) Transfection of *Plasmodium falciparum* within human red blood cells Proc Natl Acad Sci U S A 92, 973–977

87. Rosario, V. (1981) Cloning of naturally occurring mixed infections of malaria parasites Science 212, 1037–1038

88. Howieson, V. M., Tran, E., Hoegl, A., Fam, H. L., Fu, J., Sivonen, K. et al. (2016) Triazole substitution of a labile amide bond stabilizes pantothenamides and improves their antiplasmodial potency Antimicrob Agents Chemother 60, 7146–7152

89. Saliba, K. J., Horner, H. A., and Kirk, K. (1998) Transport and metabolism of the essential vitamin pantothenic acid in human erythrocytes infected with the malaria parasite Plasmodium falciparum J Biol Chem 273, 10190–10195

90. Spry, C., Saliba, K. J., and Strauss, E. (2014) A miniaturized assay for measuring small molecule phosphorylation in the presence of complex matrices Anal Biochem 451, 76–78

91. Somogyi, M. (1945) Determination of Blood Sugar Journal of Biological Chemistry 160, 69–73

92. Roos, K., Wu, C., Damm, W., Reboul, M., Stevenson, J. M., Lu, C. et al. (2019) OPLS3e: Extending force field coverage for drug-like small molecules J Chem Theory Comput 15, 1863–1874

